# Reduced coenzyme Q synthesis confers non-target site resistance to the herbicide thaxtomin A

**DOI:** 10.1101/2022.09.13.507736

**Authors:** Chloe Casey, Thomas Köcher, Clément Champion, Katharina Jandrasits, Magdalena Mosiolek, Clémence Bonnot, Liam Dolan

## Abstract

Herbicide resistance in weeds is a growing threat to global crop production. Non-target site resistance is problematic because a single resistance allele can confer tolerance to many herbicides (cross resistance), and it is often a polygenic trait so it can be difficult to identify the molecular mechanisms involved. Most characterized molecular mechanisms of non-target site resistance are caused by gain-of-function mutations in genes from a few key gene families – the mechanisms of resistance caused by loss-of-function mutations remain unclear. In this study, we first show that the mechanism of non-target site resistance to the herbicide thaxtomin A conferred by loss-of-function of the gene *PAM16* is conserved in *Marchantia polymorpha*, validating its use as a model species with which to study non-target site resistance. To identify mechanisms of non-target site resistance caused by loss-of-function mutations, we generated 10^7^ UV-B mutagenized *M. polymorpha* spores and screened for resistance to the herbicide thaxtomin A. We isolated 13 thaxtomin A-resistant mutants and found that 3 mutants carried candidate resistance-conferring SNPs in the Mp*RTN4IP1L* gene. Mp*rtn4ip1l* mutants are defective in coenzyme Q biosynthesis and accumulate higher levels of reactive oxygen species (ROS) than wild-type plants. Mutants are also defective in thaxtomin A metabolism, consistent with the hypothesis that loss of Mp*RTN4IP1L* function confers non-target site resistance. We conclude that loss of Mp*RTN4IP1L* function is a novel mechanism of non-target site herbicide resistance, and propose that other mutations which increase ROS levels or decrease thaxtomin A metabolism could confer thaxtomin A resistance in the field.

**AUTHOR SUMMARY:** Modern agriculture relies on herbicides to control weed populations. However, herbicide resistance in weeds threatens the efficacy of herbicides and global crop production, similar to how antibiotic resistance poses a global health threat. Understanding the molecular mechanisms behind herbicide resistance helps to prevent resistance from evolving and to better manage herbicide resistant weeds in the field. Here, we use a forward genetic approach in the model species *Marchantia polymorpha* to discover novel mechanisms of herbicide resistance. We report the discovery of a novel mechanism of herbicide resistance caused by loss-of-function mutations in the Mp*RTN4IP1L* gene. We find that Mp*rtn4ip1l* mutants are resistant to the herbicides thaxtomin A and isoxaben, accumulate higher levels of reactive oxygen species than wild type plants, and are defective in thaxtomin A metabolism. We predict that loss-of-function mutations or treatments that increase reactive oxygen species production could contribute to thaxtomin A tolerance.

## INTRODUCTION

The control of weeds by herbicides is threatened by the evolution of resistance in the field (1, 2). This is analogous to how human and animal health is threatened by the evolution of antibiotic resistance in bacteria. The intense selection pressure resulting from herbicide treatments can lead to the rapid evolution of herbicide resistance in the agricultural landscape (1). Resistance can result from genetic changes in the gene encoding the herbicide target which lead to conformational changes in the target protein blocking herbicide binding, or overexpression of the target (3, 4). This form of resistance, which involves mutations only in the gene encoding the target protein, is known as target site resistance. Resistance can also result from genetic changes in a variety of genes which prevent inhibitory levels of herbicide reaching the target, or that alleviate the toxic effects of the herbicide downstream from the target (1, 3, 5). For example, a mutation which causes constitutive expression of a glycolsyltransferase gene causes resistance in the weed species *Alopecurus myosuroides*. This is due to increased herbicide metabolism and increased accumulation of antioxidant flavonol metabolites which confer resistance to herbicides whose toxicity depends on the accumulation of toxic reactive oxygen species (6–8). These types of resistance – which do not involve mutations in the gene encoding the herbicide target – are known as non-target site resistance.

Both target site and non-target site herbicide resistance can be caused by gain of function mutations, which cause an increase in expression of a gene or enhanced activity of a protein. For example, EPSPS encodes a protein essential for aromatic amino acid biosynthesis which is the target of the herbicide glyphosate (9). Gain of function mutations in the gene encoding 5-enolpyruvyl shikiminate-3-phosphate synthase (EPSPS) resulting from gene amplification (multiple rounds of gene duplication) confers target site resistance to the herbicide glyphosate (10, 11). EPSPS gene copy number is greater in many glyphosate-resistant populations than in glyphosate-sensitive populations, resulting in higher steady state levels of EPSPS mRNA, protein, and total enzyme activity. The resulting elevated levels of EPSPS activity increase glyphosate resistance (11). Non-target site resistance can also be conferred by spontaneous gain of function mutations in genes encoding enzyme activities that chemically modify or compartmentalize herbicides and prevent the herbicide reaching the target (1, 5, 12). For example, constitutive overexpression of an ABC transporter causing glyphosate export into the apoplast results in glyphosate resistance in *Echinocloa colona* (13).

Loss-of-function mutations – mutations that result in reduced gene and protein function – can also cause target site resistance. This is because mutations in the gene encoding a herbicide target which cause conformational changes that prevent herbicide binding can also reduce the capacity of the protein to carry out its function. For example, some mutations in EPSPS that confer glyphosate resistance by preventing the binding of glyphosate also reduce the affinity of EPSPS for its substrate phosphoenolpyruvate, leading to reduced catalytic activity (14). However, very few loss-of-function mutations causing non-target site resistance have been reported in weeds. Recent genome wide association studies have identified mutations associated with glyphosate resistance in the weed species *A. tuberculatus* and *I. purpurea* of which some are likely to be loss-of-function, suggesting that the contribution of loss-of-function mutations to non-target site resistance has been underestimated, however the contribution of these mutations to resistance has not yet been experimentally confirmed (15–17). Only two mechanisms of non-target site resistance involving loss-of-function mutations have been proven to confer resistance, using forward genetics in the model species *A. thaliana* (18, 19). As such, the role of loss-of-function mutations in non-target site resistance in weeds is unclear.

It is likely that both gain- and loss-of-function mutations contribute to non-target site resistance in herbicide resistant weed populations, but it is difficult to identify resistance mechanisms of non-target site resistance, including those conferred by loss of gene function. This is partly because non-target site resistance is usually a polygenic trait, and each single allele may contribute little to resistance. For example, in a sample of glyphosate resistant *Amaranthus tuberculatus* individuals, it was estimated that 25 % of the variance in resistance was attributed to non-target site resistance associated with SNPs in 274 genes with a range of allele frequencies and effect sizes (15, 17). Identifying the mutant genes that contribute such a small amount to total resistance phenotypes is challenging in weeds. This makes defining the molecular basis of non-target site resistance difficult. The few alleles which have been identified are those which have a large effect on resistance. So far, these have all been gain-of-function alleles involving genes from a few key gene families such as cytochrome P450s, glutathione-S-transferases, glycosyltransferases, and ABC transporters (5). There are likely a variety of undiscovered non-target site resistance causing mutations which each have a small effect on resistance but additively cause strong resistance. Identifying these diverse mechanisms of non-target site resistance – which likely include loss-of-function mutations – remains a challenge.

We set out to identify mechanisms of non-target site resistance to thaxtomin A – a natural product being developed as a herbicide – caused by loss-of-function mutations. We chose thaxtomin A because it has been approved by the US Environmental Protection Agency as a herbicide (EPA registration number 84059-12) (20–26). Thaxtomin A is thought to act by inhibiting plant cell wall biosynthesis, a process necessary for plant growth and the inhibition of which leads to plant death (27, 28). This plant specific mode of action makes thaxtomin A a favourable herbicide choice due to its low toxicity to animals. To date, only one mechanism of resistance to thaxtomin A has been identified (via forward mutagenesis in *Arabidopsis thaliana*) caused by loss-of-function mutations in *PAM16*, which encodes a member of the TIM complex, involved in protein transport into mitochondria (18, 29, 30).

To identify novel loss-of-function mutations that confer resistance to thaxtomin A, we UV mutagenized a large population of *Marchantia polymorpha* spores and screened for mutants which survived the lethal dose of thaxtomin A. We chose *M. polymorpha* as a model species because a single female sporophyte produces millions of haploid spores which can be mutagenized and directly screened for resistance. This enables rapid mutant screens in large populations that would be difficult to carry out at the same scale in diploid flowering plant models (31, 32). Furthermore, there is evidence that gene networks underlying bryophyte gametophyte physiology and development are similar to those in angiosperms (33, 34). We identified 13 thaxtomin A-resistant mutants. Three independent mutants carried candidate resistance-conferring mutations in the Mp*RTN4IP1L* gene, which functions in the biosynthesis of coenzyme Q, a component of the mitochondrial electron transport chain. Mp*rtn4ip1l* mutants accumulated higher levels of reactive oxygen species (ROS) than wild type controls and were defective in thaxtomin A metabolism. Taken together these data suggest that mutations that lead to ROS accumulation or which hinder thaxtomin A metabolism may confer resistance to thaxtomin A. We predict that loss-of-function mutations that increase ROS levels or decrease thaxtomin A metabolism in weeds will confer resistance to thaxtomin A in the field.

## RESULTS

### A mechanism of non-target site resistance conferred by loss-of-function of *PAM16* is conserved in *M. polymorpha*

To determine if *M. polymorpha* is a suitable model species for discovering novel mechanisms of non-target site resistance to thaxtomin A, we first tested if *M. polymorpha* is sensitive to thaxtomin A. Wild type gemmae from Takaragaike-1 (Tak-1) and Takaragaike-2 (Tak-2) accessions were grown on solid medium supplemented with different concentrations of thaxtomin A. The area of living tissue of these plants was measured after 14 days (Fig 1A). Thaxtomin A treatment inhibits the growth of wild type *M. polymorpha* in a dose-dependent manner (Fig 1A and 1B). The IC_50_ (dose at which average area of living tissue is reduced by 50 %) was 56 ± 6 nM for Tak-1, and 135 ± 24 nM for Tak-2. The LD_100_ (lethal dose at which no plant tissue is alive) for both Tak-1 and Tak-2 was 5 µM, although in rare cases Tak-1 can survive this dose (Fig 1A and 1B).

**Fig 1.**
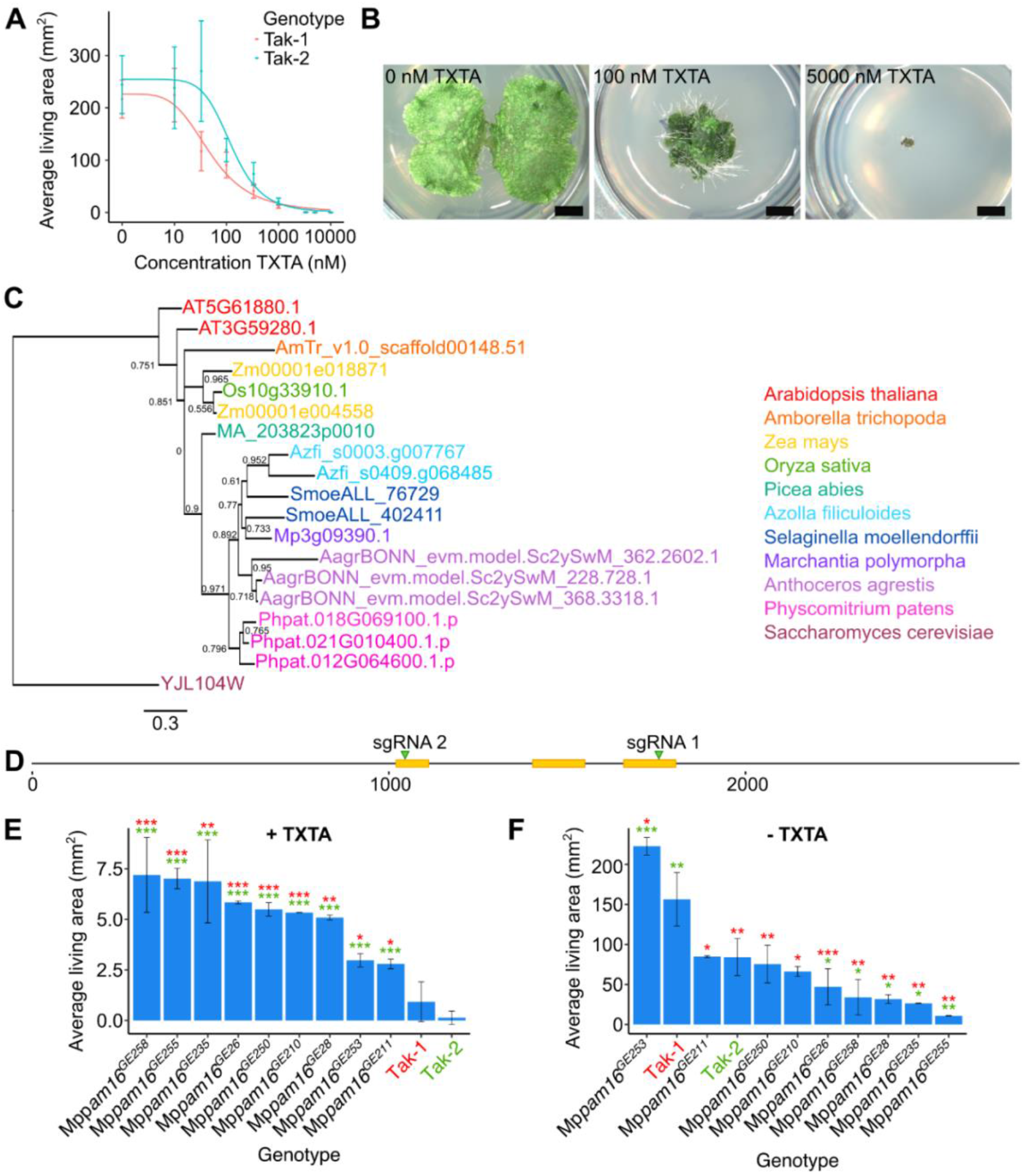
A mechanism of non-target site resistance conferred by loss-of-function of *PAM16* is conserved in *M. polymorpha*. **A:** Dose-response curves of the growth of wild type lines on thaxtomin A (TXTA). Wild type gemmae (Tak-1 and Tak-2) were grown for 14 days on solid medium supplemented with different concentrations of thaxtomin A. The fitted curves and IC_50values_ were calculated using the four-parameter log-logistic equation. Error bars represent ± standard deviation (n = 18). **B:** Wild type (Tak-1) gemmalings grown for 12 days on solid medium supplemented with DMSO, 100 nM thaxtomin A, or 5000 nM thaxtomin A. Images were taken with a Keyence VHX-7000. Scale bars represent 2 mm. **C:** Phylogenetic analysis of *PAM16* in plants. Proteins similar to AtPAM16 (At3G59280) were identified by protein BLAST search (E value < 1E-5) against the reference proteomes of various species (Table 1) (35). Orthologues were aligned and a maximum likelihood analysis was conducted. The tree was rooted with the *PAM16* homologue from *S. cerevisiae*. A non-parametric approximate likelihood ratio test based on a Shimodaira-Hasegawa-like procedure was used to calculate branch support values (36). **D:** Schematic representation of the Mp*PAM16* (Mp3g09390) gene. CDS regions are represented in yellow. The regions of the gene targeted by guide RNAs are marked with green arrowheads. **E and F:** Wild type gemmae (Tak-1 and Tak-2) and gemmae from the nine thaxtomin A-resistant Mp*pam16* mutants were grown for 12 days on solid medium supplemented with 5 µM thaxtomin A (**E**) or 0.1 % DMSO (**F**). The fitted curves and IC_50_ values were calculated using the four-parameter log-logistic equation. Error bars represent ± standard deviation (n = 2-7). Stars represent the level of significance (as determined by Student’s t-tests) of the difference between mutant and control lines (comparison to Tak-1 in red and Tak-2 in green): * = p < 0.05, ** = p < 0.01, *** = p < 0.001.

**Table 1.**
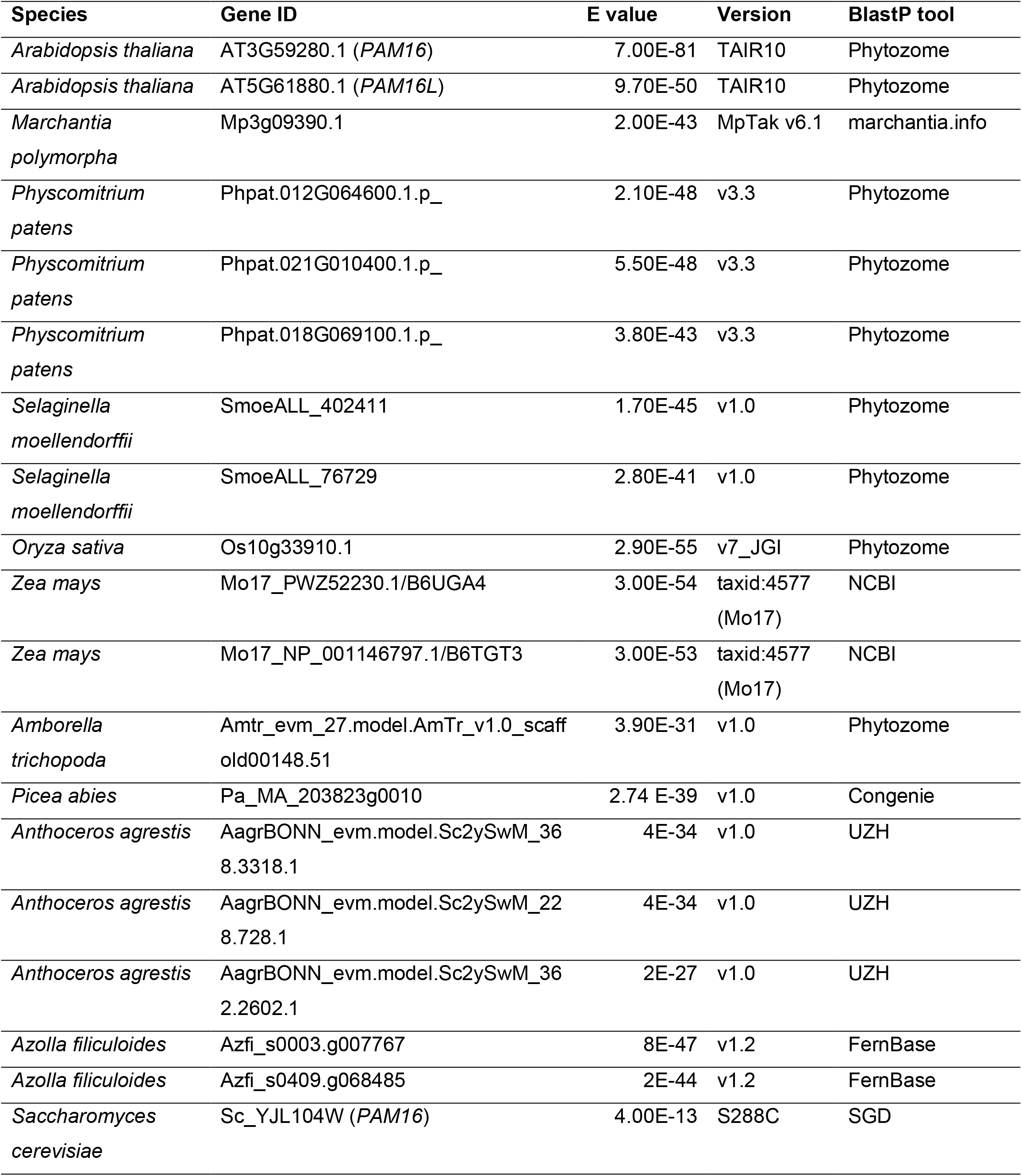
Homologues of the *A. thaliana* gene AT3G59280 identified by BlastP search (E value < 1E-5)

To validate the suitability of *M. polymorpha* to study non-target site resistance to thaxtomin A, we tested whether a known mechanism of non-target site thaxtomin A resistance discovered in *Arabidopsis thaliana* is conserved in *M. polymorpha*. Loss-of-function mutations in the At*PAM16* gene (AT3G59280) confer non-target site resistance to thaxtomin A in *A. thaliana* (18, 30). *PAM16* encodes a subunit of the PAM/TIM complex that transports proteins across the inner mitochondrial membrane (29). We tested the hypothesis that loss of *PAM16* function confers thaxtomin A-resistance in *M. polymorpha*.

The closest homologue of At*PAM16* in *M. polymorpha* was identified by BlastP as Mp3g09390 (35). To determine the evolutionary history of the *PAM16* gene and determine if Mp3g09390 is the only copy of PAM16 in *M. polymorpha*, a phylogeny was created of the homologues of *PAM16* from different species. Protein sequences similar to AtPAM16 (AT3G59280) from 11 species were identified by searching their proteomes using the BlastP algorithm (Table 1) (35). The plant species included were the liverwort *M. polymorpha*, the moss *P. patens*, the hornwort *A. agrestis*, the lycophyte *S. moellendorfii*, the fern *A. filiculoides*, the gymnosperm *P. abies*, and the angiosperms *A. trichopoda, A. thaliana*, *Z. mays*, and *O. sativa*. *PAM16* from the yeast *Saccharomyces cerevisiae* was used as an outgroup. The sequences were aligned using the L-INS-i strategy in MAFFT (37) and manually trimmed to remove poorly conserved regions (S1A and S1B Fig). A maximum likelihood tree was constructed using this alignment and a non-parametric approximate likelihood ratio test based on a Shimodaira-Hasegawa-like procedure was used to calculate node support values using PhyML 3.0 (36) (Fig 1C). There were between one and three copies of *PAM16* in each species and there was one copy in *M. polymorpha* (Mp3g09390). Mp3g09390 is therefore the only *PAM16* gene in the *M. polymorpha* genome and will be referred to as Mp*PAM16*.

To determine if loss of Mp*PAM16* function confers thaxtomin A resistance, Mp*pam16* loss-of-function mutants were generated using CRISPR-Cas9 mutagenesis. Guide RNAs were designed to generate mutations in the highly conserved N-terminal predicted signal peptide which targets the PAM16 protein to the inner mitochondrial membrane (sgRNA 2), and a non-conserved region at the C-terminal end of the protein (sgRNA 1) (Fig 1D). Thirty-three Mp*pam16* mutants were generated. The highly conserved signal peptide was mutated in 28 Mp*pam16* mutants; the non-conserved C-terminal region of the protein was mutated in the 5 remaining Mp*pam16* mutants.

Gemmae from the Mp*pam16* mutant lines were grown on solid medium supplemented with 5 µM thaxtomin A (LD_100_) (Fig 1E) or 0.1 % DMSO (Fig 1F). The area of living plant tissue was quantified after 12 days of growth. Mp*pam16* mutant lines which were significantly larger than wild type plants on thaxtomin A were classed as resistant. Nine Mp*pam16* lines were significantly larger than both Tak-1 and Tak-2 on 5 µM thaxtomin A and therefore classed as thaxtomin A-resistant (p < 0.05) (Fig 1E).

Five of the nine thaxtomin A-resistant Mp*pam16* mutants were significantly smaller than Tak-1 and Tak-2 plants in control conditions, including the four Mp*pam16* mutants which grew largest on thaxtomin A – Mp*pam16^GE258^*, Mp*pam16^GE255^*, Mp*pam16^GE235^*, and Mp*pam16^GE26^* (Fig 1F). This suggests that growth defects are more likely in Mp*pam16* mutant lines with stronger resistance to thaxtomin A.

All 9 thaxtomin A-resistant Mp*pam16* mutants were mutated in the highly conserved region that comprises the predicted PAM16 signal peptide (S1C Fig). The mutations in the thaxtomin A-resistant mutants affect between 2 and 19 amino acids and involve a range of substitutions, insertions, and deletions (S1C Fig). A mutation in the signal peptide is predicted to disrupt PAM16 function by preventing its transport to the mitochondria. These mutations are therefore likely to be loss-of-function. Given that all 9 thaxtomin A-resistant Mp*pam16* mutant lines carry likely loss-of-function mutations, we conclude that loss-of-function of Mp*pam16* can confer thaxtomin A resistance. The resistance of Mp*pam16* mutants to thaxtomin A validates *M. polymorpha* as a model to identify novel mechanisms of non-target site resistance to this herbicide.

### 13 thaxtomin A-resistant *M. polymorpha* mutants were isolated from a screen of 10^7^ mutants

A genetic screen was carried out to identify mutants resistant to the herbicide thaxtomin A. To generate thaxtomin A-resistant mutants, 2 x 10^7^ wild type *M. polymorpha* spores were plated on solid medium supplemented with 5 µM thaxtomin A and exposed to UV-B irradiation to induce mutations. A screening dose of 5 µM thaxtomin A (LD_100_) was selected as screening at the lethal dose (as opposed to higher than the lethal dose) preferentially selects for non-target site resistance (38). The dose of UV-B radiation used killed approximately 50 % of spores (S2 Fig). Ten million mutagenized spores were screened for resistance to thaxtomin A 14 days after mutagenesis (Fig 2A). Resistant mutants were transferred to fresh solid medium supplemented with 5 µM thaxtomin A to confirm their resistance. Mutants that survived the second round of selection were retained, and gemmae from each resistant mutant were plated onto fresh solid medium supplemented with 5 µM thaxtomin A to verify the retention of resistance through asexual reproduction (Fig 2B). Mutant gemmae that survived and grew significantly larger than wild type gemmae on 5 µM thaxtomin A were classed as thaxtomin A-resistant (Fig 2C). Using this protocol, 13 thaxtomin A-resistant Mp*thar* (*th*axtomin *A r*esistant) mutant lines were identified out of the 10^7^ mutagenized spores screened.

**Fig 2.**
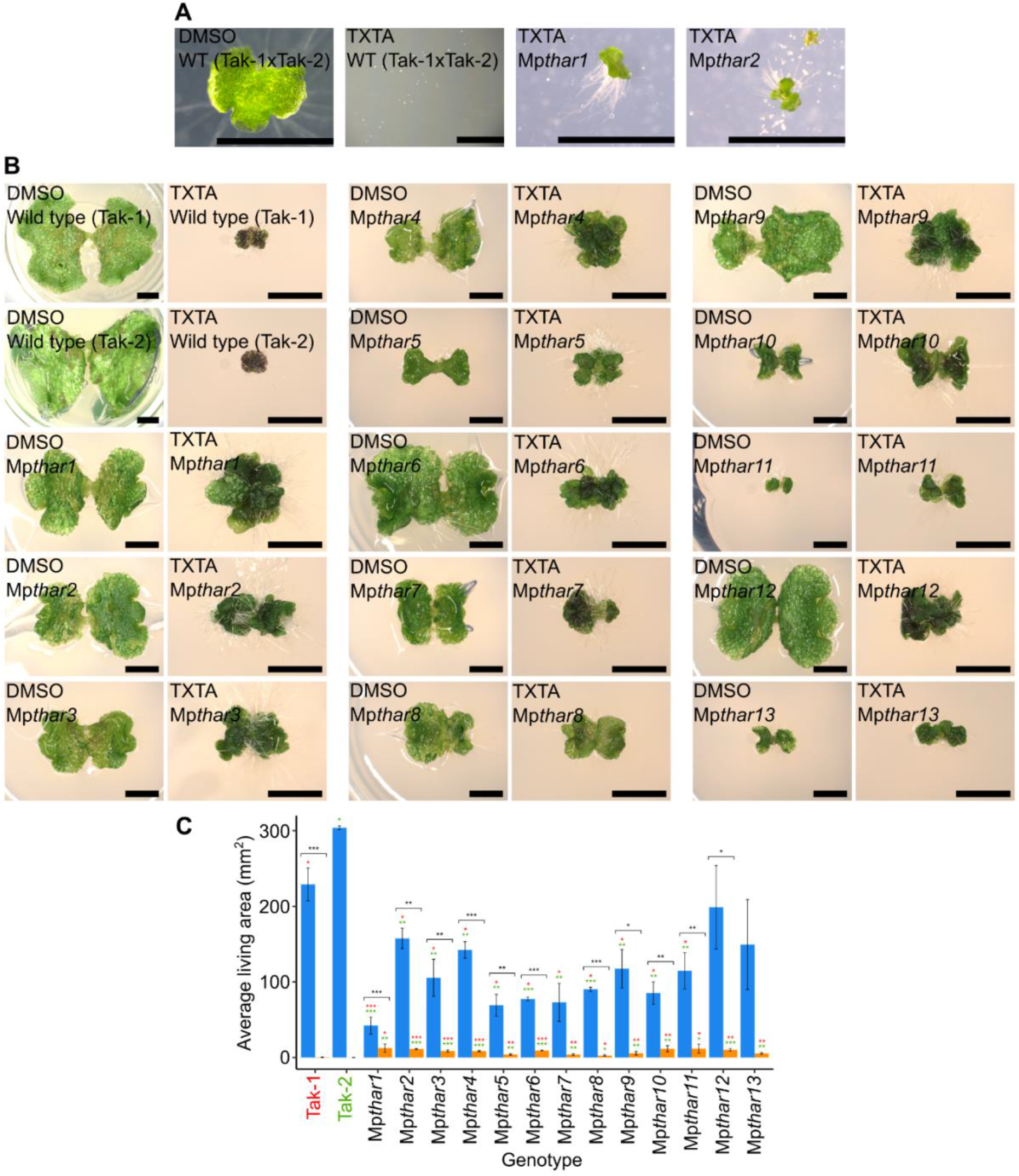
13 thaxtomin A-resistant mutants were isolated from a screen of 10^7^ mutants. **A:** Wild type spores from a cross between Tak-1 and Tak-2 grown for 14 days on solid medium supplemented with 0.1 % DMSO or 5 µM thaxtomin A (TXTA), and UV-B mutagenesis selection plates supplemented with 5 µM thaxtomin A showing 14-day old thaxtomin A-resistant mutants Mp*thar*1 and Mp*thar*2 surrounded by dead mutagenized thaxtomin A-sensitive spores. Images were taken with a Leica DFC310 FX stereomicroscope. Scale bars represent 2 mm. **B:** Wild type (Tak-1 and Tak-2) and Mp*thar* gemmalings grown for 12 days on solid medium supplemented with 0.1 % DMSO or 5 μM thaxtomin A. Images were taken with a Keyence VHX-7000 and the maximum pixel value was adjusted to 188 in ImageJ. Scale bars represent 2 mm. **C:** Gemmae from wild type lines (Tak-1 and Tak-2) and from Mp*thar* lines were plated on solid medium supplemented with DMSO (blue bars) or 5 μM thaxtomin A (orange bars) and grown for 21 days (n = 3-13). The fitted curves and IC_50_ values were calculated using the four-parameter log-logistic equation. Error bars represent ± standard deviation. Stars represent the level of significance (as determined by Student’s t-tests) of the difference between mutant and control lines subjected to the same treatment (comparison to Tak-1 in red and Tak-2 in green), or between individuals of the same genotype subjected to different treatments (black): * = p < 0.05, ** = p < 0.01, *** = p < 0.001.

### The Mp*RTN4IP1L* gene is mutated in three Mp*thar* mutants

To identify mutations that confer thaxtomin A resistance, the genome of each Mp*thar* mutant was sequenced and the most likely resistance-conferring SNP in each mutant was identified using a modified version of a SNP calling bioinformatic pipeline adapted from (39). Mismatches in the genomes of Mp*thar* mutants were compared to mismatches in the genomes of thaxtomin A-sensitive plants (Tak-1, Tak-2, and thaxtomin A-sensitive UV-B mutants (39)). Mismatches found only in Mp*thar* mutants and which passed further filtering steps were considered to be candidate resistance-conferring mutations. Between 6 and 27 candidate resistance-conferring mutations were identified for each Mp*thar* mutant. These SNPs were compared between Mp*thar* mutants. There were candidate resistance-conferring SNPs in the Mp3g19030 gene in 3 independent Mp*thar* mutants (Mp*thar2*, Mp*thar4*, and Mp*thar6*) (Fig 3A and 3B). The presence of these SNPs was confirmed by Sanger sequencing (S3A, S3B, and S3C Fig). Given that the Mp3g19030 gene carries a candidate resistance-conferring mutation in 3 independent Mp*thar* mutants, it is likely that a mutation in Mp3g19030 confers thaxtomin A resistance.

**Fig 3.**
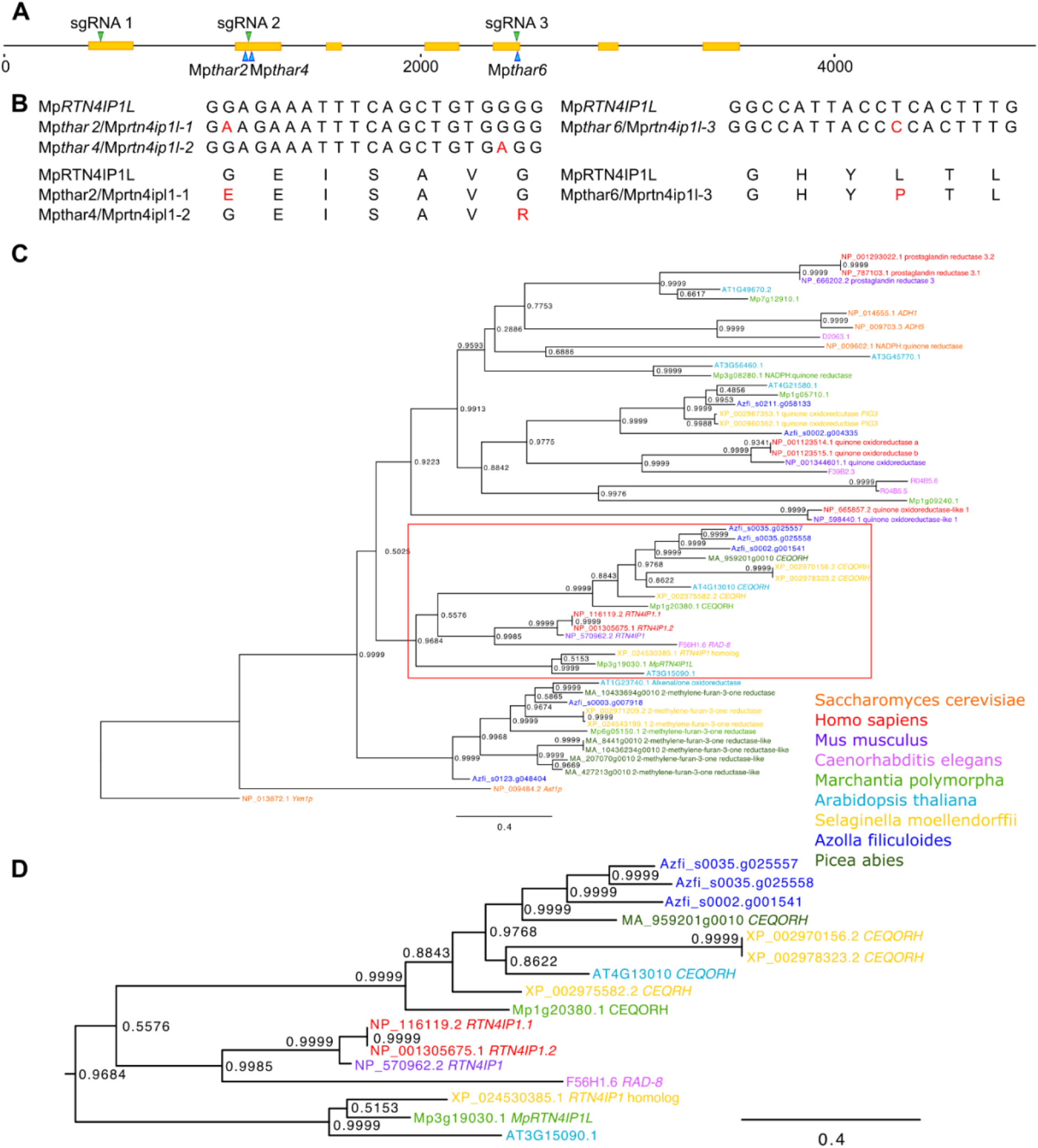
The Mp*RTN4IP1L* (Mp3g19030) gene is mutated in 3 Mp*thar* mutants. **A:** Schematic representation of the Mp*RTN4IP1L* (Mp3g19030) gene. CDS regions are represented in yellow. The positions of the SNPs in Mp*thar* are marked with blue arrowheads. The regions of the gene targeted by guide RNAs (sgRNA 1 and sgRNA 2) are marked with green arrowheads. **B:** Nucleotide alignments of the wild type Mp*RTN4IP1L* gene and the mutant Mp*thar* genes, and amino acid alignments of the MpRTN4IP1L protein and predicted mutant Mp*thar* proteins. **C:** Phylogenetic analysis of *RTN4IP1* in eukaryotes. Proteins similar to the human RTN4IP1 protein were identified by protein BLAST search (E value < 1E-5) against the reference proteomes of various species (Table 2) (35). Orthologues were aligned and a maximum likelihood analysis was conducted. The tree was rooted with the *RTN4IP1* homologue (Yim1p) from *S. cerevisiae*. A non-parametric approximate likelihood ratio test based on a Shimodaira-Hasegawa-like procedure was used to calculate branch support values (36). **D:** Magnification of the red boxed branch from (**C**) showing that Mp3g19030 diverges into a monophyletic clade containing the human *RTN4IP1* gene.

**Table 2.** Species included in the phylogenetic analysis of Hs*RTN4IP1.1* homologues. Proteins similar to HsRTN4IP1.1 were identified via BlastP search of proteomes from these species (35). Only homologues with an E value < 1 E-5 were included in the analysis.

| Organism | BLAST tool | Number of *RTN4IP1* homologues (E < 1E-5) |
| --- | --- | --- |
| *Homo sapiens* | NCBI | 7 |
| *Saccharomyces cerevisiae* | NCBI | 5 |
| *Caenorhabditis elegans* | Wormbase | 5 |
| *Mus musculus* | NCBI | 4 |
| *Marchantia polymorpha* | marchantia.info | 7 |
| *Arabidopsis thaliana* | TAIR | 7 |
| *Selaginella moellendorffii* | NCBI | 8 |
| *Picea abies* | Congenie | 6 |
| *Azolla filiculoides* | Fernbase | 7 |

Mp3g19030 is annotated in the reference *M. polymorpha* genome (MpTak v6.1) as a homologue of the *H. sapiens RETICULON 4-INTERACTING PROTEIN 1.1* (Hs*RTN4IP1.1*) gene. Hs*RTN4IP1.1* encodes a mitochondrial NADH oxidoreductase involved in the biosynthesis of Coenzyme Q, and its homologue *RAD-8* was first discovered in *C. elegans* (40, 41). To determine the phylogenetic relationship between Mp3g19030 and its homologues, protein sequences similar to HsRTN4IP1.1 from 9 species were identified using the BlastP algorithm to search their proteomes (35). These species included *H. sapiens*, *S. cerevisiae*, *M. musculus*, *C. elegans*, *M. polymorpha*, *A. thaliana*, *S. moellendorffii*, *A. filiculoides*, and *P. abies* (Table 2). The protein sequences were aligned via the L-INS-i strategy in MAFFT (37) and trimmed to remove poorly conserved regions (S3D Fig). A maximum likelihood tree was constructed and a non-parametric approximate likelihood ratio test based on a Shimodaira-Hasegawa-like procedure was used to calculate node support values using PhyML 3.0 (36). The *RTN4IP1* homologue (*Yim1p*) from the yeast *S. cerevisiae* was used to root the tree (Fig 3C).

There were between 4 and 8 Hs*RTN4IP1.1* homologues in each of the species analyzed (Table 2). The homologues group into a clade of 2-methylene-furan-3-one reductases (support value 0.9999), a clade containing Hs*RTN4IP1.1* and its closest homologues (support value 0.9684), and a clade containing quinone oxidoreductases, alcohol dehydrogenases, and prostaglandin reductases (support value 0.9223) (Fig 3C). Based on the annotations, the identified homologues of Hs*RTN4IP1.1* therefore likely include enzymes with similar activities.

Mp3g19030 forms a clade (support value 0.9999) with the closest Hs*RTN4IP1.1* homologues from *A. thaliana* and *S. moellendorffii* (AT3G15090.1; E value 6 x 10^-54^, and XP_024530385.1; E value 1 x 10^-46^) that is sister to the clade representing the *RAD-8/RTN4IP1* genes from *H. sapiens*, *M. musculus* (*RTN4IP1*; E value 0) and *C. elegans* (*RAD-8*; E value 4 x 10^-63^) (support value 0.9985) (Fig 3D). Mp3g19030 is the most closely related *M. polymorpha* homologue of Hs*RTN4IP1.1* gene based on BlastP. However, the topology of the phylogenetic tree suggests that Mp1g20380, a chloroplast envelope quinone oxidoreductase homologue, is the closest *M. polymorpha* homologue of Hs*RTN4IP1.1* (Fig 3D). Therefore, Mp3g19030 will be referred to as Mp*RTN4IP1-LIKE* (Mp*RTN4IP1L*).

### Loss-of-function of Mp*RTN4IP1L* is a novel mechanism of herbicide resistance

To independently verify that loss of Mp*RTN4IP1L* function confers thaxtomin A resistance, Mp*rtn4ip1l* loss-of-function mutants were generated using CRISPR-Cas9 targeted mutagenesis. A guide RNA was designed to mutate the beginning of the gene to induce frameshifts (sgRNA 1), and 2 guide RNAs were designed to mutate the sites mutated in Mp*thar*2, Mp*thar*4, and Mp*thar*6 (sgRNA 2 and sgRNA 3) (Fig 3A). In total, 19 Mp*rtn4ip1l* mutants were generated. Based on the nucleotide and predicted protein sequences, there were in frame indels in Mp*rtn4ip1l* which affected 5 or fewer amino acids in the Mprtn4ip1l protein of Mp*rtn4ip1l^GE111^* and Mp*rtn4ip1l^GE117^* (S4 Fig). There were large indels or frameshift mutations in Mp*rtn4ip1l* which affected 10 or more amino acids in the Mprtn4ip1l protein of the remaining 17 Mp*rtn4ip1l* mutants, therefore these 17 lines were predicted loss-of-function lines (S4 Fig).

Gemmae from the Mp*rtn4ip1l* mutant lines and from lines with a wild type Mp*RTN4IP1L* (Tak-1, Tak-2, and 3 lines which were transformed with the CRISPR construct but had no mutations in the Mp*RTN4IP1L* gene; Mp*RTN4IP1L^GE153^*, Mp*RTN4IP1L^GE314^*, and Mp*RTN4IP1L*^GE356^) were grown on solid medium supplemented with 5 µM thaxtomin A (LD_100_) (Fig 4A and 4B) or 0.1 % DMSO (Fig 4A and 4C). The average area of living plant tissue was quantified after 12 days of growth. Plant lines which grew significantly larger than Tak-1 and Tak-2 plants on thaxtomin A were classed as resistant.

**Fig 4.**
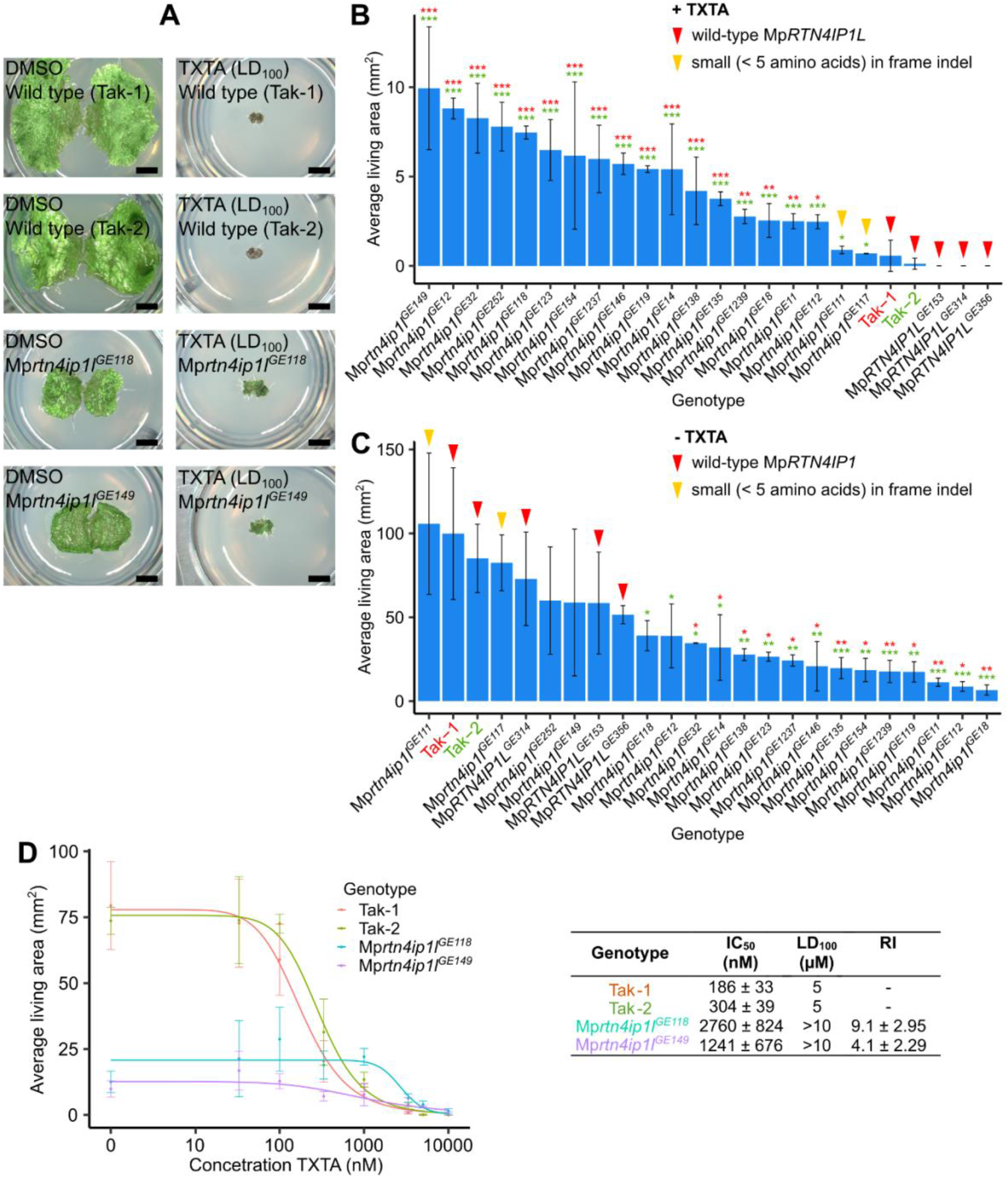
Loss-of-function of Mp*RTN4IP1L* confers resistance to thaxtomin A. **A:** Gemmae from wild type lines (Tak-1 and Tak-2) and from Mp*rtn4ip1l* mutants (Mp*rtn4ip1l*^GE118^ and Mp*rtn4ip1l*^GE149^) were grown for 12 days on solid medium supplemented with DMSO or 5 μM thaxtomin A (TXTA). Images were taken with a Keyence VHX-7000 and the maximum pixel value was adjusted to 200 in ImageJ. **B and C:** Wild type gemmae (Tak-1 and Tak-2) and gemmae from Mp*rtn4ip1l* mutants were grown for 12 days on solid medium supplemented with 5 µM thaxtomin A (**B**) or 0.1 % DMSO (**C**). Error bars represent ± standard deviation (n = 2-6). Stars represent the level of significance (as determined by Student’s t-tests) of the difference between mutant and control lines (comparison to Tak-1 in red and Tak-2 in green): * = p < 0.05, ** = p < 0.01, *** = p < 0.001. Red arrowheads indicate lines with a wild type copy of Mp*RTN4IP1L*; yellow arrowheads indicate lines with a small (< 5 amino acids) in frame mutation in Mp*RTN4IP1L*. **D:** Dose-response curves of the growth of wild type and Mp*rtn4ip1l* mutants on thaxtomin A, and table of corresponding IC_50_, LD_100_, and RI values. Gemmae from wild type lines (Tak-1 and Tak-2) and from two independent Mp*rtn4ip1l* mutants generated via CRISPR-Cas9 mutagenesis (Mp*rtn4ip1l*^GE118^ and Mp*rtn4ip1l*^GE149^) were grown for 12 days on solid medium supplemented with different concentrations of thaxtomin A. The fitted curves and IC_50_ values were calculated using the four-parameter log-logistic equation. Error bars represent ± standard deviation (n = 3).

The 17 predicted loss-of-function Mp*rtn4ip1l* lines were significantly larger than both Tak-1 and Tak-2 on 5 µM thaxtomin A so were classed as thaxtomin A-resistant (p<0.05) (Fig 4B). The 2 mutants with small in frame indels in Mp*rtn4ip1l* and the lines with a wild type copy of Mp*RTN4IP1L* however did not grow significantly larger than Tak-1 and Tak-2 on 5 µM thaxtomin A so were classed as thaxtomin A-sensitive (Fig 4B). This suggests that loss-of-function of Mp*RTN4IP1L* confers resistance to thaxtomin A.

When grown in control conditions, the areas of plant lines with a wild type copy of Mp*RTN4IP1L* and the 2 Mp*rtn4ip1l* mutants with small in frame indels are statistically indistinguishable from wild type (Fig 4C). However, 13 of the 17 predicted loss-of-function Mp*rtn4ip1l* mutants were significantly smaller than wild type lines in control conditions (Fig 4C). Loss-of-function of Mp*RTN4IP1L* therefore likely causes decreased growth in control conditions.

To quantify the thaxtomin A resistance of Mp*rtn4ip1l* CRISPR mutants, gemmae from 2 independent Mp*rtn4ip1l* CRISPR lines (Mp*rtn4ipl1l*^GE118^ and Mp*rtn4ip1l*^GE149^) were grown on solid medium supplemented with different doses of thaxtomin A (Fig 4A and 4D). The average area of living plant tissue was quantified after 12 days of growth. Dose-response curves were fitted using the four-parameter log-logistic equation (Fig 4D). Three parameters of resistance were calculated from the curves: IC_50_ (the concentration of thaxtomin A at which the area of living tissue is reduced by 50 %), LD_100_ (lethal dose at which there is no surviving tissue), and RI (resistance index; ratio between wild type and mutant IC_50_). The IC_50_ of Tak-2 was used to calculate the most conservative estimate of RI of each Mp*rtn4ip1l* line. The IC_50_, RI, and LD_100_ values of both Mp*rtn4ip1l* mutant lines were significantly higher than either wild type line (Fig 4D). Based on their RI, Mp*rtn4ip1l*^GE1518^ and Mp*rad8*^GE1549^ are respectively at least 9.1 ± 2.95 times and 4.1 ± 2.29 times more resistant to thaxtomin A than wild type (Fig 4D). Taken together, these data indicate that loss-of-function mutations in Mp*RTN4IP1L* confer thaxtomin A resistance.

### Mp*rtn4ip1l* mutants produce lower levels of coenzyme Q and accumulate higher levels of reactive oxygen species than wild type plants

Having shown that loss-of-function mutations in the Mp*RTN4IP1L* gene confer thaxtomin A resistance, we sought to determine how these mutations changed the physiology of plants to make them thaxtomin A resistant. HsRTN4IP1 interacts with proteins involved in coenzyme Q (CoQ) synthesis and knock-out mutants in the mouse *RTN4IP1* gene accumulate less CoQ than wild type (41). We therefore hypothesized that CoQ levels would be lower in Mp*rtn4ip1l* mutants than in wild type *M. polymorpha*. To test this hypothesis, we measured the levels of coenzyme Q10 (CoQ) in the extracts of wild type (Tak-1) and Mp*rtn4ip1l^GE149^*mutants via LC-MS/MS. The levels of CoQ in Mp*rtn4ip1l^GE149^* are significantly lower than wild type levels in both control and TXTA-treated conditions, consistent with the hypothesis that Mp*RTN4IP1L* is involved in CoQ synthesis (Fig 5A).

**Fig 5.**
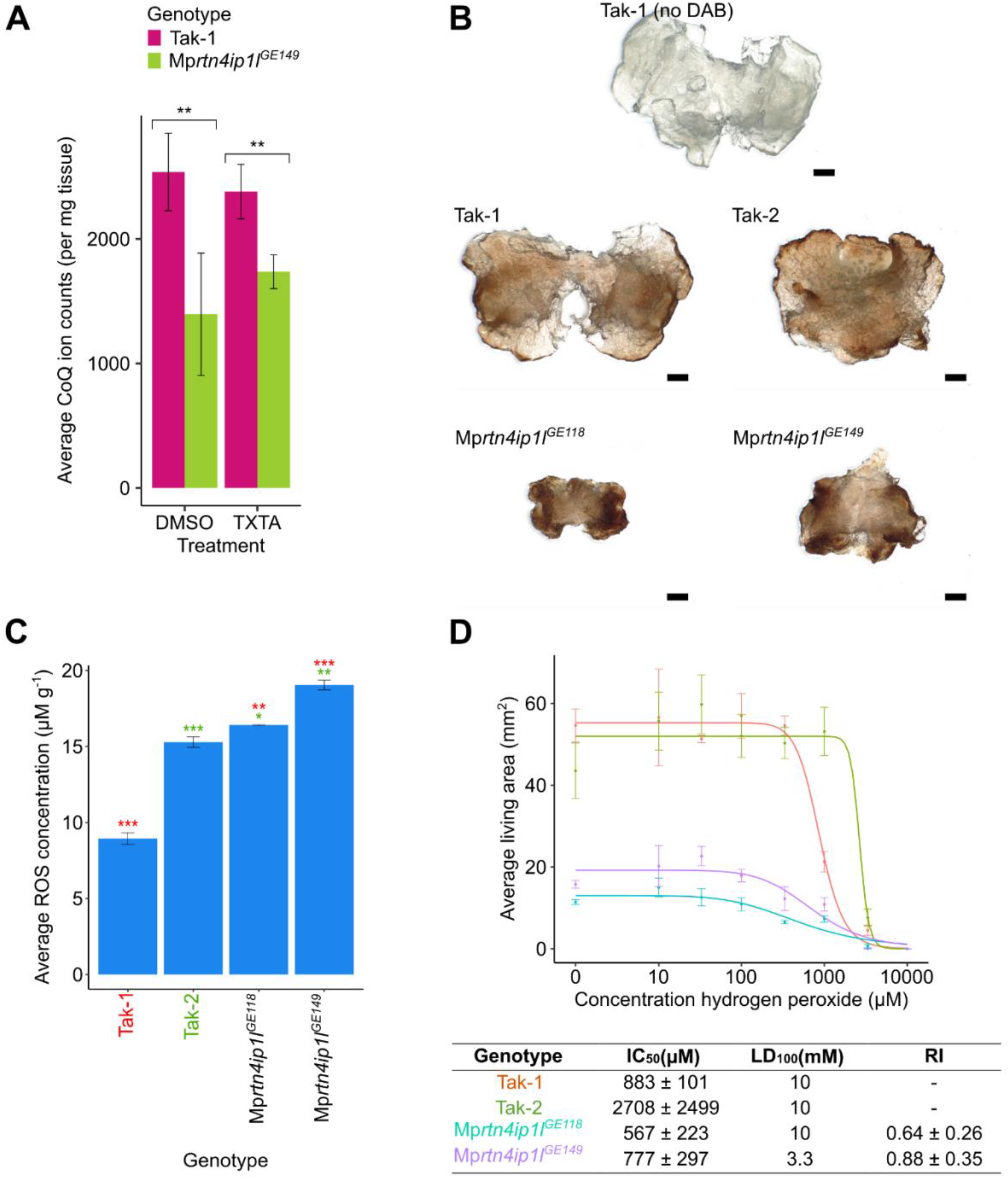
Mp*rtn4ip1l* mutants are defective in coenzyme Q synthesis and overproduce reactive oxygen species. **A:** Ion counts of coenzyme Q10 (CoQ) in wild-type and Mp*rtn4ip1l* mutants as quantified by targeted LC-MS/MS. Tak-1 and Mp*rtn4ip1l^GE149^* gemmae were grown on solid medium supplemented with 0.1 % DMSO for 14 days, then transferred to solid medium supplemented with 0.1% DMSO or 5 μM thaxtomin A for 2 days. Cellular fractions were extracted from the plants and analysed via LC-MS/MS. Stars represent the level of significance of the difference between Tak-1 and Mp*rtn4ip1l^GE149^* samples as determined by Student’s t-tests: * = p < 0.05, ** = p < 0.01, *** = p < 0.001. Error bars represent ± standard deviation (n = 3-6). **B:** DAB staining of Tak-1, Tak-2, Mp*rtn4ip1l*^GE118^, and Mp*rtn4ip1l*^GE149^. 12-day old gemmalings were stained with 3,3’-diaminobenzidine (DAB), which forms a brown precipitate upon reaction with H_2_O_2_. Stained plants were bleached to remove chlorophyll and imaged with a Keyence VHX-7000. Scale bars represent 500 μm. **C:** Concentration of ROS in wild type and Mp*rtn4ip1l* mutants. The concentration of ROS in 12-day old Tak-1, Tak-2, Mp*rtn4ip1l*^GE118^, and Mp*rtn4ip1l*^GE149^ plants was determined by homogenising frozen samples in perchloric acid, followed by incubation with ferric xylenol-orange (FOX) and measurement of absorbance at 560 nm using an Ultrospec 3100 pro spectrophotometer. The absorbance measurements were converted to concentrations of ROS using a calibration curve. Stars represent the level of significance of the difference between mutant and control lines (Tak-1 in red and Tak-2 in green) as determined by Student’s t-tests: * = p < 0.05, ** = p < 0.01, *** = p < 0.001. Error bars represent ± standard deviation (n=3). **D:** Dose-response curves of the growth of wild type and Mp*rtn4ip1l* mutants on H_2_O_2_. Gemmae from Tak-1 (orange), Tak-2 (green), Mp*rtn4ip1l*^GE118^ (blue), and Mp*rtn4ip1l*^GE149^ (purple) were grown for 10 days on solid medium supplemented with different concentrations of H_2_O_2_. The fitted curves and IC_50_ values were calculated using the four-parameter log-logistic equation. Error bars represent ± standard deviation (n = 3). LD_100_ is the lowest concentration at which area of all replicates was 0. RI is the ratio between each Mp*rtn4ip1l* mutant’s IC_50_ and the IC_50_ of Tak-1, the more sensitive wild type line, to calculate the most conservative estimate of RI.

Coenzyme Q is an electron acceptor that transports electrons from Complexes I and II to Complex III of the oxidative phosphorylation electron transport chain (42). As electrons flow through the electron transport chain, electrons may leak from Complexes I, II and III and react with oxygen to form reactive oxygen species (ROS). The binding site of CoQ at Complex I is one of the sites of ROS production in the mitochondria (42). Reduced levels of CoQ result in fewer electron carriers active in the oxidative phosphorylation transport chain leading to an excess of electrons leaking from the chain and reacting with oxygen to form ROS. Furthermore, non-mitochondrial CoQ acts as a lipid soluble antioxidant, protecting against oxidative stress by reacting with lipid peroxide radicals thereby preventing their ability to cause lipid peroxidation (43–45). Therefore, we hypothesized that ROS levels would be higher in Mp*rtn4ip1l* mutants than in wild type plants.

To test the hypothesis that Mp*rtn4ip1l* mutants produce higher levels of ROS than wild type plants, 12-day old gemmalings of Tak-1, Tak-2, Mp*rtn4ip1l*^GE118^ and Mp*rtn4ip1l*^GE149^ were stained with 3,3’-diaminobenzidine (DAB). DAB reacts with hydrogen peroxide (H_2_O_2_) – a form of ROS – to form a brown precipitate. The amount of brown precipitate is proportional to the amount of H_2_O_2_ in each sample (46). Both Mp*rtn4ip1l* mutants stained darker than the two wild type lines, indicating higher levels of H_2_O_2_ in mutant thalli than in wild type (Fig 5B). The concentration of various forms of ROS (including H_2_O_2_ and lipid hydroperoxides) was also quantified in Mp*rtn4ip1l* mutants using a modified ferric-xylenol orange (FOX) assay (47, 48). ROS concentrations were significantly higher in Mp*rtn4ip1l*^GE118^ and Mp*rtn4ip1l*^GE149^ than in wild type plants (Fig 5C).

Since Mp*rtn4ip1l* mutants produce more ROS than wild type, we hypothesized that Mp*rtn4ip1l* mutants would be more sensitive (hypersensitive) to ROS treatment than wild type plants. To test this hypothesis the sensitivity of Mp*rtn4ip1l* mutants to toxic levels of exogenous ROS was compared to that of wild type plants. Tak-1, Tak-2, Mp*rtn4ip1l*^GE118^ and Mp*rtn4ip1l*^GE149^ gemmae were grown on solid medium supplemented with a range of H_2_O_2_ concentrations. The average area of living plant tissue was quantified after 10 days of growth. Dose-response curves were fitted using the four-parameter log-logistic equation, and resistance parameters IC_50_, LD_100_ and RI were calculated from the curves (Fig 5D). Both Mp*rtn4ip1l* mutants were more sensitive to H_2_O_2_ than wild type lines. The RI values of Mp*rtn4ip1l*^GE118^ and Mp*rtn4ip1l*^GE149^ were 0.64 ± 0.26 and 0.88 ± 0.35 respectively, demonstrating that these lines are more sensitive to H_2_O_2_ than either wild type line. The RI value of Mp*rtn4ip1l*^GE118^ was statistically significant (Fig 5D). While the RI value of Mp*rtn4ip1l*^GE1549^ was not statistically significant, the LD_100_ of Mp*rtn4ip1l*^GE149^ was lower than either wild type line (Fig 5D). These data demonstrate that Mp*rtn4ip1l* mutants are more sensitive to H_2_O_2_ than wild type plants.

Together, these data indicate that ROS levels are higher in Mp*rtn4ip1l* mutants than in wild type plants. Addition of H_2_O_2_ to *A. thaliana* cells in culture confers partial thaxtomin A resistance (49). The stiffening of the cell wall observed upon H_2_O_2_ treatment is thought to prevent the toxic action of thaxtomin A on cell wall biosynthesis (49). We hypothesize that the reduced levels of CoQ in Mp*rtn4ip1l* mutants causes thaxtomin A resistance by increasing ROS levels which changes the physiology of the plant making thaxtomin A less effective. According to this hypothesis, loss-of-function of Mp*rtn4ip1l* is a mechanism of non-target site resistance.

### Loss-of-function of Mp*RTN4IP1L* likely confers non-target site resistance

To test the hypothesis that loss-of-function of *RTN4IP1L* results in non-target site resistance, the strength of thaxtomin A resistance and the cross resistance to other herbicides were tested in Mp*rtn4ip1l* mutants. Herbicide cross resistance and weak thaxtomin A resistance would be consistent with the hypothesis that *RTN4IP1L* loss-of-function mutations confer non-target site resistance.

The strength of thaxtomin A resistance in Mp*rtn4ip1l* mutants was measured and compared to the strength of resistance to chlorsulfuron in chlorsulfuron target-site resistant mutants. Chlorsulfuron is a herbicide which targets the enzyme acetohydroxyacid synthase (AHAS), also known as acetolactate synthase (ALS). We generated 5 *M. polymorpha* chlorsulfuron-resistant target site resistant mutants – Mpahas^*chlorsulfuron resistant*^ (Mp*ahas^chlr^*) – in a UV-B mutagenesis screen for resistance to chlorsulfuron. Each mutant carries a target site resistance mutation in Mp*AHAS* which confers chlorsulfuron resistance (S6 Fig). Mutations causing a Pro197Ser or Pro197Leu amino acid change, which have been observed to confer chlorsulfuron resistance in angiosperm weeds, were present in 4 out of the 5 Mp*ahas^chlr^* mutants isolated in this screen (2, 50, 51).

To compare the strength of herbicide resistance in Mp*rtn4ip1l* and Mp*ahas^chlr^* lines, gemmae from 2 independent Mp*rtn4ip1l* mutants (Mp*rtn4ip1l*^GE118^ and Mp*rtn4ip1l*^GE149^) were grown on solid medium supplemented with LD_100_ or 5 x LD_100_ of thaxtomin A (Fig 6A and 6B), and gemmae from 5 independent Mp*ahas^chlr^* mutant lines were grown on solid medium supplemented with LD_100_ or 5 x LD_100_ of CS (Fig 6C and 6D). The average area of living plant tissue was quantified after 12 days of growth.

**Fig 6.**
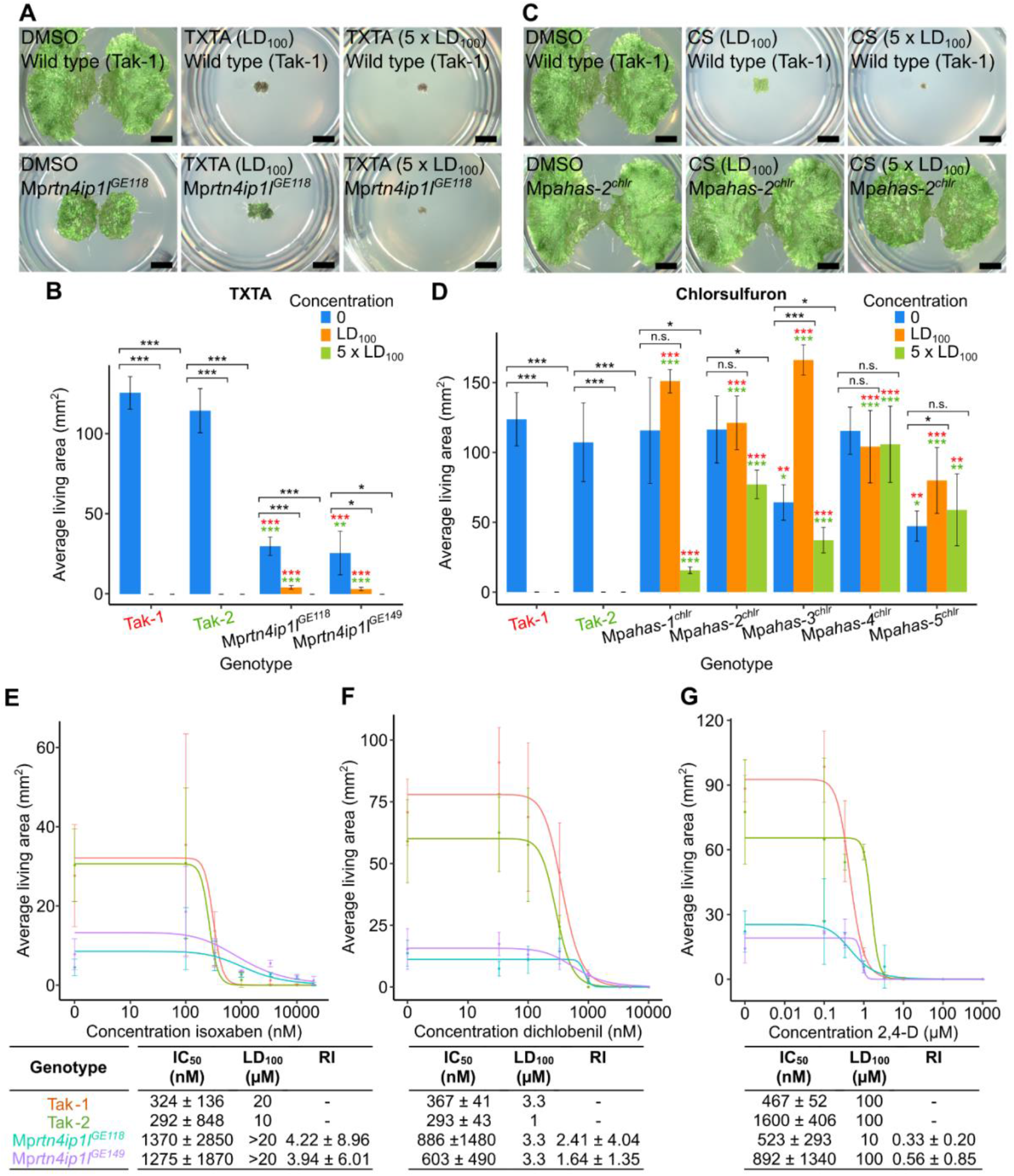
Loss-of-function of Mp*RTN4IP1L* likely confers non-target site resistance. **A and B:** Wild type gemmae (Tak-1 and Tak-2) and gemmae from Mp*rtn4ip1l* mutants were grown on solid medium supplemented with LD_100_ (5 µM) or 5 x LD_100_ (25 µM) of thaxtomin A. **(A)** Gemmalings were imaged using a Keyence VHX-7000 (scale bars represent 2 mm) **(B)** Error bars represent ± standard deviation (n = 3). Stars represent the level of significance (as determined by Student’s t-tests) of the difference between mutant and control lines (comparison to Tak-1 in red and Tak-2 in green), or between individuals of the same genotype subjected to different treatments (black): * = p < 0.05, ** = p < 0.01, *** = p < 0.001. **C and D:** Wild type gemmae (Tak-1 and Tak-2) and gemmae from Mp*ahas^chlr^* mutants were grown on solid medium supplemented with LD_100_ (33 nM) or 5 x LD_100_ (165 nM) of chlorsulfuron. **(C)** Gemmalings were imaged using a Keyence VHX-7000 (scale bars represent 2 mm) **(D)** Error bars represent ± standard deviation (n = 3). Stars represent the level of significance (as determined by Student’s t-tests) of the difference between mutant and control lines (comparison to Tak-1 in red and Tak-2 in green), or between individuals of the same genotype subjected to different treatments (black): * = p < 0.05, ** = p < 0.01, *** = p < 0.001. **E,F, and G:** Dose-response curves of the growth of Mp*rtn4ip1l* mutants on herbicides with different targets and tables of corresponding IC_50_, LD_100_, and RI values. Gemmae from Tak-1 (orange), Tak-2 (green), Mp*rtn4ip1l^GE118^*(blue), and Mp*rtn4ip1l^GE149^* (purple) were grown for 12 days on solid medium supplemented with different doses of isoxaben (**E**), dichlobenil (**F**), or 2,4-D (**G**). The fitted curves and IC_50_ values were calculated using the four-parameter log-logistic equation. Error bars represent ± standard deviation (n = 3). LD_100_ is the lowest concentration at which area of all replicates was 0. RI was calculated as the ratio between each Mp*rtn4ip1l* mutant’s IC_50_ and the IC_50_ of the most resistant wild type line to calculate the most conservative estimate of RI.

Mp*rtn4ip1l* mutants are significantly larger than wild type plants on the thaxtomin A LD_100_, confirming their thaxtomin A resistance (Fig 6A and 6B). Similarly, Mp*ahas^chlr^* mutants are significantly larger than wild type on the chlorsulfuron LD_100_ (Fig 6C and 6D). However, Mp*rtn4ip1l* mutants grow significantly less on thaxtomin A LD_100_ than in control conditions (Fig 6A and 6B). Conversely, Mp*ahas*^chlr^ mutants are either statistically indistinguishable or significantly larger when grown on chlorsulfuron LD_100_ than in control conditions (Fig 6C and 6D). This suggests that Mp*rtn4ip1l* mutants are weakly resistant to thaxtomin A, whereas Mp*ahas^chlr^* are strongly resistant to chlorsulfuron. This is consistent with the hypothesis that Mp*rtn4ip1l* mutants are non-target site resistant.

Mp*rtn4ip1l* mutants die when grown on five times the LD_100_ of thaxtomin A (Fig 6A and 6B), whereas Mp*ahas*^chlr^ mutants do not die on five times the LD_100_ of chlorsulfuron (Fig 6C and 6D). The inability of Mp*rtn4ip1l* mutants to survive 5 x LD_100_ is also consistent with the hypothesis that Mp*rtn4ip1l* mutants are weakly resistant to thaxtomin A. Resistance from loss-of-function mutations in Mp*rtn4ip1l* is therefore more likely to be non-target site resistance than target site resistance.

To further test the hypothesis that loss-of-function of Mp*rtn4ip1l* function causes non-target site resistance against thaxtomin A, we tested if Mp*rtn4ip1l* mutants were resistant to other herbicides (cross resistance). To quantify the cross resistance of Mp*rtn4ip1l* mutants to different herbicides, the resistance of Mp*rtn4ip1l^GE118^*and Mp*rtn4ip1l^GE149^* to the herbicides isoxaben, dichlobenil, and 2,4-D was tested. Isoxaben targets cellulose biosynthesis, targeting the cellulose synthase subunits CESA1, CESA3, and CESA6 (52). Dichlobenil is also classed as a cellulose biosynthesis inhibitor, but its target protein is still unknown (53). 2,4-D is an auxin mimic, causing deregulation of the auxin signalling pathway (54). Gemmae from Tak-1, Tak-2, Mp*rtn4ip1l^GE118^*, and Mp*rtn4ip1l^GE149^* were grown on solid medium supplemented with different doses of isoxaben (Fig 6E), dichlobenil (Fig 6F), or 2,4-D (Fig 6G). After 10 days of growth, the area of living tissue was quantified, and dose-response curves were fitted using the four-parameter log-logistic equation. IC_50_, LD_100_, and RI for wild type and Mp*rtn4ip1l* mutant lines were calculated from the resulting dose-response curves (Fig 6E, 6F, and 6G).

Mp*rtn4ip1l* mutant lines were resistant to isoxaben, Mp*rtn4ip1l^GE118^* with an RI of 4.22 ± 8.96 and Mp*rtn4ip1l^GE149^* with an RI of 3.94 ± 6.01. Although these RI values are not statistically significant, the LD_100_ of both Mp*rtn4ip1l* mutant lines was higher than wild type; both were alive when grown on 20 µM isoxaben which is lethal to Tak-1 and Tak-2 (Fig 6E). These data demonstrate that Mp*rtn4ip1l^GE118^* and Mp*rtn4ip1l^GE149^* are resistant to isoxaben. The cross resistance of Mp*rtn4ip1l* mutants to isoxaben is consistent with the hypothesis that loss of Mp*RTN4IP1L* function confers non-target site resistance.

Mp*rtn4ip1l* mutant lines were not resistant to dichlobenil or 2,4-D. The RI values of Mp*rtn4ip1l^GE118^* and Mp*rtn4ip1l^GE149^* on dichlobenil are not statistically significant, and neither mutant can survive at a higher concentration of dichlobenil than wild type (Fig 6F). Mp*rtn4ip1l^GE118^* has an RI value of 0.33 ± 0.20 on 2,4-D so is significantly 2,4-D sensitive, while the RI of Mp*rtn4ip1l^GE149^*on 2,4-D is not statistically significant. Neither Mp*rtn4ip1l* mutant has a higher LD_100_ than wild type on 2,4-D (Fig 6G). Mp*rtn4ip1l* lines are therefore not cross resistant to dichlobenil or 2,4-D.

Taken together, the weak thaxtomin A resistance and isoxaben cross resistance of Mp*rtn4ip1l* mutants demonstrate that loss-of-function of Mp*RTN4IP1L* is likely to confer non-target site resistance. Non-target site resistance is often caused by chemical modification of a herbicide to a non-toxic product, or by reduced herbicide uptake (1, 5). Non-target site resistance to thaxtomin A in Mp*rtn4ip1l* mutants could be caused by a change in herbicide metabolism or uptake that result from elevated ROS levels in Mp*rtn4ip1l* mutants caused by defective CoQ synthesis.

### Mp*rtn4ip1l* mutants are defective in thaxtomin A metabolism

We set out to test the hypothesis that non-target site resistance in Mpr*tn4ip1l* mutants results from differences in the metabolism or uptake of thaxtomin A between wild type and mutants. To test this hypothesis, thaxtomin A and putative thaxtomin A metabolites in thaxtomin A-treated samples of Tak-2 wild type and Mp*rtn4ip1l*^GE118^ mutants were detected via a precursor ion analysis for the two most abundant fragment ions of thaxtomin A using LC/MS-MS (Fig 7A and 7B). There is a peak of pure thaxtomin A (retention time – RT – 8.99 min) and a peak of an unknown compound (RT 9.25 min) in the chromatogram of wild type Tak-2 (Fig 7A). Both peaks are absent in the analysis of the untreated samples (S7A Fig). The two prominent fragment ions of thaxtomin A are present in the MS/MS spectra of the unknown compound, suggesting that it is a thaxtomin A derivative (S7B and S7C Fig). The molecular mass of the unknown compound is 45 Da less than pure thaxtomin A. An explanation for these observations is that the unknown compound is a derivative of thaxtomin A that lacks the NO_2_ group (delta mass of 45 Da) (Fig 7C and 7D). This suggests that wild type *M. polymorpha* metabolizes thaxtomin A by removing the NO_2_ group of the 4-nitro-tryptophan moiety. These two peaks are present in the chromatogram from Mp*rtn4ip1l*^GE149^. However, the RT 9.25 peak is considerably smaller in Mp*rtn4ip1l*^GE149^ than wild type (Fig 7B). This suggests that Mp*rtn4ip1l* mutants accumulate less of the derivative and are therefore defective in thaxtomin A metabolism.

**Fig 7.**
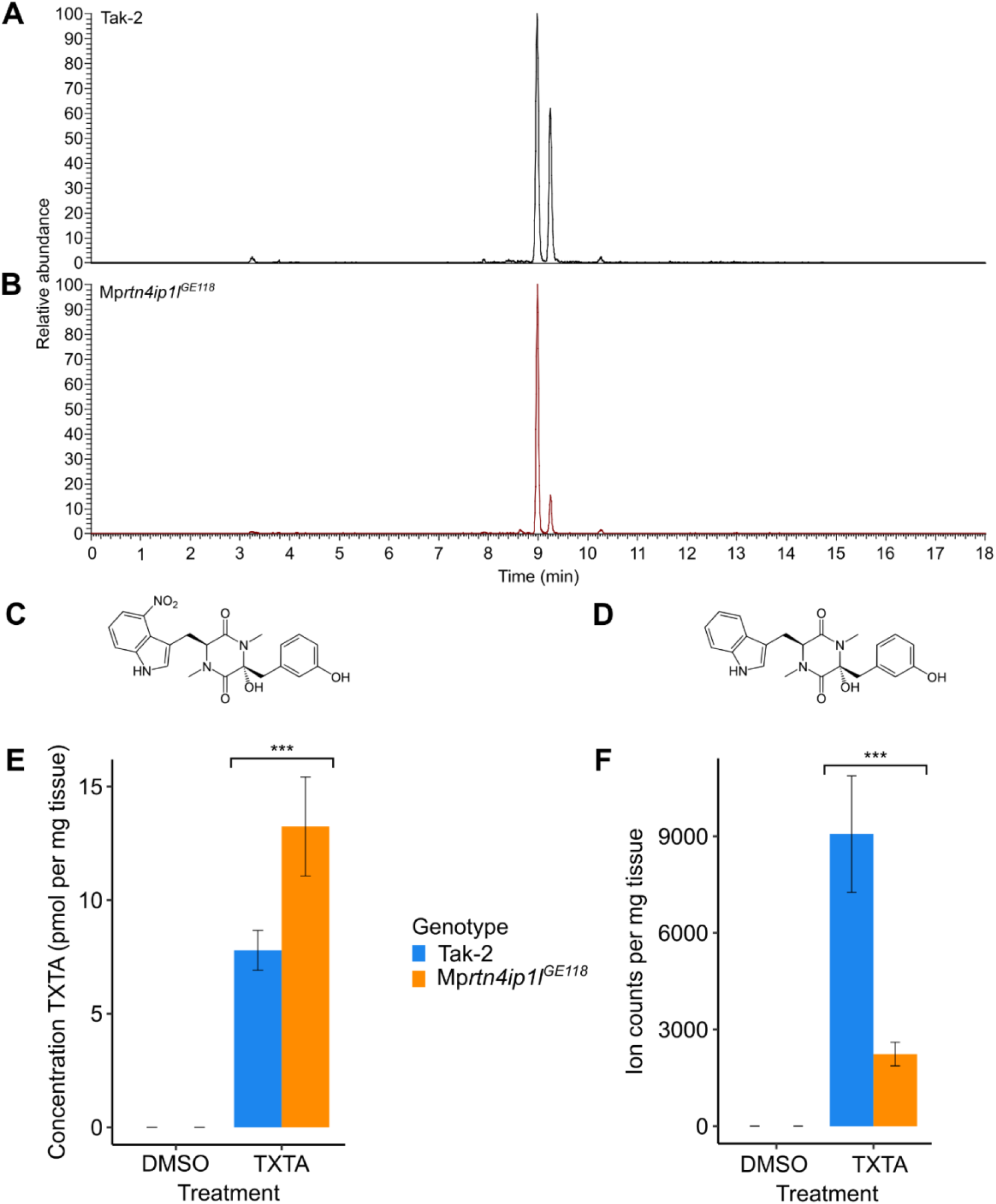
Mp*rtn4ip1l* mutants are defective in thaxtomin A metabolism. **A and B:** LC-MS/MS analysis of thaxtomin A and its putative metabolite in Tak-2 **(A)** and Mp*rtn4ip1l*^GE118^ **(B)** extracts. Gemmae from each line were grown on solid medium supplemented with 0.1 % DMSO for 14 days, then transferred to solid 1 medium supplemented with 5 μM thaxtomin A and grown for 2 days. Cellular fractions were extracted from thaxtomin A-treated samples and a precursor ion analysis was conducted via LC-MS/MS (n = 6). Chromatograms of the total ion currents of precursor ion scanning for *m/z* 247.1 are depicted. The peak eluting at a retention time of 8.99 min. is thaxtomin A (*m/z* 439.1), the peak eluting at a retention time of 9.25 (*m/z* 394.1) is its putative metabolite. **C:** Molecular structure of thaxtomin A **D:** Molecular structure of putative thaxtomin A metabolite **E:** Concentration of pure thaxtomin A (TXTA) in DMSO and herbicide-treated samples of Tak-2 and Mp*rtn4ip1l*^GE118^ as quantified by targeted LC/MS-MS. **F:** Ion counts (normalised by weight) of the putative thaxtomin A metabolite in DMSO and herbicide-treated samples of Tak-2 and Mp*rtn4ip1l*^GE118^ as quantified by targeted LC-MS/MS.

To quantify the potential defect in thaxtomin A metabolism in Mp*rtn4ip1l* mutants, we measured the relative amounts of thaxtomin A (RT 8.99) and the putative thaxtomin A derivative (RT 9.25 min) in wild type and mutant plants using targeted LC-MS/MS employing selected reaction monitoring (Fig 7E and 7F). The concentration of thaxtomin A is significantly higher in Mp*rtn4ip1l*^GE118^ than in Tak-2 (p < 0.05) (Fig 7E). This is consistent with the hypothesis that the chemical modification of thaxtomin A is defective in the mutants (Fig 7E). Furthermore, this is inconsistent with the hypothesis that Mp*rtn4ip1l* mutants uptake less thaxtomin A than wild type plants, suggesting that their resistance is not conferred by reduced herbicide uptake.

The ion counts of the putative thaxtomin A derivative (RT 9.25) are significantly lower in Mp*rtn4ip1l*^GE118^ than in Tak-2 (p < 0.05), consistent with the hypothesis that the chemical modification of thaxtomin A is defective in the mutants (Fig 7F). Although the difference in the concentration of thaxtomin A between the samples cannot be directly compared to the difference in ion counts of the unknown compound, these data suggest that wild type plants metabolize thaxtomin A to form a derivative compound, but this metabolism is defective in Mp*rtn4ip1l* mutants. Given that Mp*rtn4ip1l* mutants are resistant to thaxtomin A, this is consistent with the hypothesis that thaxtomin A metabolism has a toxic effect on the plant.

To independently verify that the metabolism of thaxtomin A is defective in Mp*rtn4ip1l* mutants, the presence of thaxtomin A and its putative metabolite were also detected via precursor ion analysis in Tak-1 wild type and Mp*rtn4ip1l*^GE149^ mutants. The peak corresponding to the putative metabolite was stronger in Tak-1 than in Mp*rtn4ip1*^GE149^, supporting the hypothesis that thaxtomin A metabolism is defective in Mp*rtn4ip1* mutants (S7D Fig).

Together these data indicate that a thaxtomin A derivative that accumulates in wild type plants accumulates at much lower levels in the resistant Mp*rtn4ip1l* mutants. It is hypothesized that the metabolism of thaxtomin A is toxic to plants. According to this hypothesis, Mp*rtn4ip1l* mutants are resistant to thaxtomin A because they are defective in thaxtomin A metabolism. The high levels of ROS produced in Mp*rtn4ip1l* mutants because of low levels of CoQ may inhibit the metabolism of thaxtomin A.

## DISCUSSION

We report the discovery of a novel mechanism of non-target site resistance caused by loss-of-function mutations in the Mp*RTN4IP1L* gene using a forward genetic screen. We demonstrate that Mp*rtn4ip1l* mutants are not only resistant to thaxtomin A and isoxaben, but are also defective in thaxtomin A metabolism. Our findings demonstrate the potential of our approach to identify novel mutations conferring non-target site herbicide resistance which have been difficult to identify and characterize to date.

We propose that resistance to thaxtomin A in Mp*rtn4ip1l* mutants is the result of defective metabolism of the herbicide. Loss of Mp*RTN4IP1L* function may directly or indirectly cause the defect in thaxtomin A metabolism. One hypothesis is that the chemically modified thaxtomin A derivative is herbicidally active (bioactive) whereas unmodified thaxtomin A is not toxic. Decreased thaxtomin A metabolism in Mp*rtn4ip1l* mutants would therefore prevent the production of bioactive molecules, leading to thaxtomin A resistance. The difference in molecular mass between thaxtomin A and its derivative is 45 Da suggesting that the derivative lacks an -NO_2_ group. If true, it is consistent with the hypothesis that denitration of thaxtomin A produces the bioactive form of the herbicide. However, it has been suggested that the nitro group of thaxtomin A is required for phytotoxicity (20), and other nitroaromatic compounds are known for their toxicity (55, 56), consistent with the hypothesis that unmodified thaxtomin A is toxic. An alternative hypothesis is that the metabolism of thaxtomin A results in the formation of toxic by-products. The reduction of nitro groups in nitroaromatic compounds produces toxic nitrogen-containing radicals (57). Therefore, if the removal of the nitro group during metabolism involves its reduction, toxic nitro radicals could be formed. The nitro group may alternatively react to form other nitro group-containing toxic compounds. According to this hypothesis, the decreased thaxtomin A metabolism in Mp*rtn4ip1l* mutants would lead to decreased levels of toxic by-products, and therefore resistance to thaxtomin A. These two alternative mechanisms could explain the thaxtomin A resistance observed in Mp*rtn4ip1l* mutants. Further analysis of the chemical structure, toxicity, and production in planta of the putative thaxtomin A metabolite are necessary to elucidate its potential role in thaxtomin A toxicity.

Mp*rtn4ip1l* mutants are defective in the biosynthesis of coenzyme Q, an electron transporter in oxidative phosphorylation. Consequently, mutants produce more ROS than wild type plants. ROS are highly reactive, oxygen-containing radicals with one or more unpaired electrons. High levels of ROS cause peroxidation of cell membrane lipids leading to cell death (58, 59). Several herbicides such as paraquat or glufosinate kill plants by inducing ROS overproduction (59–61). However, at lower concentrations, ROS contribute to plant defenses, both by causing physiological changes such as cell wall strengthening, or by acting as signaling molecules (62–65). ROS pretreatment confers resistance to thaxtomin A in *A. thaliana* (49). The higher ROS levels of Mp*rtn4ip1l* mutants could therefore contribute to non-target site resistance to thaxtomin A. One hypothesis is that increased ROS levels inhibit the metabolism of thaxtomin A in Mp*rtn4ip1l* mutants leading to decreased production of toxic by-products.

Cross resistance with isoxaben suggests that resistance conferred by the Mp*rtn4ip1l* mutations constitutes non-target site resistance. However, Mp*rtn4ip1l* mutants are not resistant to dichlobenil. Thaxtomin A, isoxaben, and dichlobenil are cellulose biosynthesis inhibitors. Based on their effects on the movement and presence of cellulose synthase enzymes at the plasma membrane, thaxtomin A and isoxaben are classed as group 1 cellulose biosynthesis inhibitors and dichlobenil is classed as group 2 (28). ROS can increase the rigidity of the cell wall, which may hinder the effect of cellulose biosynthesis inhibitors (49, 62). An alternative hypothesis to explain the contribution ROS to resistance in Mp*rtn4ip1l* mutants – which takes into account their cross resistance pattern – is that the increased ROS levels alter the cell wall in a way which confers resistance to group 1 but not group 2 cellulose biosynthesis inhibitors.

The mechanism of non-target site resistance reported here is also likely to operate in angiosperm weeds and not be restricted to *M. polymorpha* or other liverworts. First, the *RTN4IP1L* gene is found throughout the angiosperms and therefore agricultural weeds, where mutation would likely result in weak resistance. Second, our data indicate that loss-of-function mutations in the *PAM16* gene confer weak thaxtomin A resistance in both *M. polymorpha* and *A. thaliana* (18). Therefore, non-target site resistance caused by loss of *PAM16* function in *M. polymorpha* also operates in angiosperms. Furthermore, we report that four chlorsulfuron-resistant *M. polymorpha* UV-B mutants carry Pro197Ser or Pro197Leu mutations, which are among the most frequently reported mechanisms of chlorsulfuron resistance observed in chlorsulfuron-resistant angiosperm weeds (2, 50, 51). Together, these data demonstrate that at least some herbicide resistance mechanisms are conserved between *M. polymorpha* and angiosperms, validating the use of *M. polymorpha* as a model with which to identify novel mechanisms of non-target site resistance which could evolve in weeds.

Mp*rtn4ip1l* mutants are smaller than wild type plants in control conditions, suggesting that loss of Mp*RTN4IP1L* function would incur a considerable fitness cost in the wild. Given this, the evolution of Mp*rtn4ip1l* loss-of-function based resistance in the field is uncertain. However, some alleles conferring herbicide resistance – both target site and non-target site – incur a fitness cost but are nevertheless maintained in weed populations due to the intense selection imposed by herbicide treatment (66–69). We propose that loss-of-function of *RTN4IP1L*, or mutations which affect the same molecular processes, could therefore evolve in weed populations treated with thaxtomin A. If loss-of-function of *RTN4IP1L* is identified in resistant weeds in the field, our findings can inform weed management practices. Based on our characterization of phenotypes of Mp*rtn4ip1l* loss-of-function mutants, weeds with *RTN4IP1L* loss-of-function mutations cannot be controlled by isoxaben. However, our data suggest that weeds that evolve thaxtomin A resistance through loss-of-function mutations in the *RTN4IP1L* genes, or through the overproduction of ROS, would be hypersensitive to herbicides which kill plants by ROS overproduction such as paraquat or glufosinate (59–61). We conclude that our methodology can uncover novel mechanisms of non-target site resistance caused by loss-of-function mutations which allows for more informed and efficient weed management in practice.

## MATERIALS AND METHODS

### Plant lines and growth conditions

Wild-type plants were *M. polymorpha* laboratory accessions Takaragaike-1 (Tak-1) and Takaragaike-2 (Tak-2) (70). Mp*pam16* and Mp*rtn4ip1l* lines were generated via CRISPR-Cas9 mutagenesis of wild-type *M. polymorpha* spores (from a cross between Tak-1 and Tak-2). All lines were maintained via asexual propagation of gemmae or thallus excision in the case of mutants which did not produce gemmae. Plants were grown on solid ½ Gamborg medium. All plant material used for experiments was grown in a growth chamber at 23 °C under 24-hour 10-30 µmol m^-2^ s^-1^ white light. Plant material to be stored long-term was kept in a growth chamber at 17 °C under 6 hours 10-30 µmol m^-2^ s^-1^ white light and 18 hours dark. To produce spores, plants were grown on a mixture of John Innes n. 2 compost and fine vermiculite at a ratio of 2:1. Plants were grown in growth chambers at 23 °C and under 16 hours white light and continuous far-red light to induce transition to the reproductive phase. Once mature, water was pipetted onto the antheridia produced by male plants to collect sperm, then transferred to archegonia produced by female plants. Sporangia were harvested before bursting and stored fresh at 4 °C in sterile dH_2_O.

### Fresh spore sterilization

Fresh sporangia were sterilized with 1 % NaDCC (sodium dichloroisocyanurate) for 4 minutes followed by rinsing with sterile dH2O.

### Chemicals and stock solution preparation

Thaxtomin A (TXTA) (SML1456), dichlobenil (45431), isoxaben (36138), chlorsulfuron (34322), 2,4-dichlorophenoxyacetic acid (2,4-D) (31518), were obtained from Sigma-Aldrich. Stock solutions were prepared by dissolving in pure dimethyl sulfoxide (DMSO) (D8418) from Sigma-Aldrich.

### Gemmaling dose-response assays

Gemmae were grown on solid ½ Gamborg medium supplemented with different concentrations of herbicide dissolved in DMSO. Gemmalings were imaged using a Berthold Nightowl II LB 983 *In Vivo* Imaging System (Berthold, Bad Wildbad, Germany). Images were taken after exposing gemmalings to 120 s white light. The imaging system detects chlorophyll autofluorescence (560 nm). The lateral area of autofluorescing (living) tissue was determined using the indiGo™ software package (Berthold, Bad Wildbad, Germany). Dose-response curves were generated using the ggplot2 and drc packages in R (71, 72); the four-parameter log-logistic equation with a fixed lower limit of 0 was used to fit a dose-response curve to the data.

### Phylogenetic analysis of PAM16 and RTN4IP1 homologues

Homologues of the *A. thaliana* At3G59280 protein or of the *H. sapiens* RTN4IP1.1 protein were identified by protein BLAST search against the reference proteomes of various species (35). Only homologues with an E value less than 1E-5 were used to construct the trees. Homologues were aligned via the L-INS-i strategy using MAFFT version 7 (37). The alignments were manually trimmed using BioEdit7.2. A maximum likelihood tree was constructed with PhyML 3.0 using an estimated gamma distribution parameter and the LG model of amino acid substitution (36). A non-parametric approximate likelihood ratio test based on a Shimodaira-Hasegawa-like procedure was used to calculate branch support values using PhyML 3.0 (36). The trees were visualized in FigTree v1.4.4 and rooted with the respective *Saccharomyces cerevisiae* homologues (PAM16 or Yim1p).

### CRISPR-Cas9 mutagenesis

CRISPR-Cas9 mutants were generated based on the protocols described in (73) and (74). Guide RNAs (sgRNAs) annealing to different parts of the genes Mp*PAM16* (Mp3g09390) or Mp*RTN4IP1L* (Mp3g19030) were designed to be 5’ of an “NGG” site (PAM sequence) as required by the CRISPR-Cas9 system (73). Guide RNAs were cloned into the pHB453 or pMpGE_En03 constructs (74) to generate the entry vectors using primers “sgRNA” (Table 3). The expression vectors were generated by LR reaction of the destination vector pMpGE010 and the entry vectors. Vectors were transformed into *Escherichia coli* One Shot OmniMAX 2 T1^R^ (Thermo Fisher Scientific: Cat. # C854003).

The expression vectors were transformed into *Agrobacterium tumefaciens* strain GV3101. Wild-type *M. polymorpha* spores from a cross between Tak-1 and Tak-2 accessions were transformed as per the protocol in (75). Transformants were selected by growth on solid ½ Gamborg medium supplemented with 10 mg/l hygromycin B (Melford: CAS# 31282-04-9) to select for plants with the CRISPR-Cas9 construct insertion, and 100 mg/l cefotaxime (Melford: CAS# 64485-93-4) to kill any remaining *A. tumefaciens*.

To identify potential mutations in the Mp*PAM16* or Mp*RTN4IP1L* genes, genomic regions including the loci targeted by the guide RNAs were amplified by PCR and Sanger sequenced using primers “PAM16_G” or “RTN4IP1L_G” (Table 3). Sanger sequencing of the purified PCR products was conducted by Source BioScience or by the Molecular Biology service at Vienna BioCenter Core Facilities (VBCF), member of the Vienna BioCenter (VBC), Austria. Mutants were named according to the gene mutated (Mp*pam16* or Mp*rtn4ip1l*), followed by “GE” (gene edited), the number of the guide RNA targeting the locus in which the mutation is found, and the number transformant that was genotyped (e.g. Mp*rtn4ip1l^GE149^*– a gene edited transformant with a mutation in the *RTN4IP1L* gene at the locus targeted by sgRNA 1 which was the 49^th^ transformant genotyped).

### Generation of herbicide-resistant *M. polymorpha* lines via UV-B mutagenesis

Spores were mutagenized according to the protocol in (39). Sterilised fresh wild type *M. polymorpha* spores were plated on solid modified Johnson’s medium supplemented with thaxtomin A (5 µM – LD_100_) or chlorsulfuron (140 nM – 4 x LD_100_). The spores were exposed to UV-B irradiation (302 nm) for an exposure time corresponding to or near the LD_50_ (an exposure time of UV-B at which 50 % of spores are killed – S2 Fig) using a UVP BioDoc-It™. Plates of mutagenized spores were wrapped in aluminium foil and left in the dark overnight. Plates were then unwrapped and placed in a Sanyo growth chamber at 23 °C in 24-hour light. After 14 days of growth, plates were screened for surviving sporelings. Surviving sporelings were transferred to solid modified Johnson’s medium supplemented with fresh herbicide. Plants which survived this second transfer onto a lethal dose of herbicide were maintained. To test for retention of resistance through asexual reproduction, gemmae from these plants were plated onto a lethal dose of herbicide. Lines whose resistance was inherited were classified as herbicide resistant lines and maintained. Lines with thaxtomin A resistance were named Mp*thar* (*<u>th</u>*axtomin *<u>A r</u>*esistant).

Chlorsulfuron-resistant lines were tested for target site resistance by PCR amplification and Sanger sequencing of regions of the MpAHAS gene using primers GCS_Fw and GCS_Rv (Table 3). Sanger sequencing of the purified PCR products was carried out by Source BioScience. Lines with target site resistance to chlorsulfuron were named Mp*ahas^chlr^* (Mp*<u>ahas chl</u>*orsulfuron *r*esistant).

### Genomic DNA sequencing

Wild-type (Tak-1, Tak-2, OxTak1F, and OxTak2M) lines and herbicide-resistant lines (Mp*thar* and Mp*ahas^chlr^*) were grown on solid ½ Gamborg medium for three weeks on in a growth chamber at 23 °C under 24-hour 10-30 µmol m^-2^ s^-1^ white light. After three weeks of growth, plant material was harvested and flash frozen in liquid nitrogen. Samples were ground in liquid nitrogen and genomic DNA was extracted using the Qiagen DNeasy Plant Maxi kit (Cat. # 68163) according to the kit protocol. After elution, the gDNA was cleaned up and concentrated using the Zymo Genomic DNA Clean & Concentrator kit (Cat. # D4010). The gDNA was eluted in nuclease-free water.

The concentration of the gDNA was checked using a Nanodrop™ 1000 spectrophotometer and a Qubit® 2.0 fluorometer according to the instruction manuals. The gDNA quality was checked by running 2 µl DNA (approximately 10 µg µl^-1^) on a 0.7 % agarose gel at 70 V for 45 minutes and checking for degradation. Genomic DNA samples which had passed quality control checks were sent for sequencing to the Next Generation Sequencing Facility at Vienna BioCenter Core Facilities (VBCF), member of the Vienna BioCenter (VBC), Austria. DNA Libraries were prepared using the Westburg NGS Library Prep kit and fragment size was determined using a BioLabTech Fragment Analyzer™. The DNA was sequenced on an Illumina NovaSeq 6000 SP flowcell using 150 bp paired-end reads.

### Non-allelism based SNP discovery pipeline

The “non-allelism based SNP discovery” pipeline from (39) was used with slight adaptations to identify candidate resistance-conferring SNPs in herbicide-resistant lines. Raw reads were trimmed to remove low-quality reads and NEB adaptors using Trimmomatic 0.38 (76). The read coverage was then normalized using khmer 2.1.2 (77). Reads were aligned to the *Marchantia polymorpha* reference genome (MpTak1 v5.1 plus the female chromosome from v3.1; now known as MpTak v6.1) using bowtie2. Reads were sorted by position and reads from different lanes were merged using samtools 1.10. The variant call analysis was carried out using samtools and bcftools (ploidy defined as haploid, multiallelic/rare variant call option selected) to generate bcf files listing mismatches between the sequencing reads and the reference genome for each line (78). For each bcf file, only mismatches which were supported by 7 – 100 reads and where the number of reads supporting high quality alternative alleles ≥ the number of reads supporting high quality reference alleles were retained (DP4[2]+DP4[3])/sum(DP4)>0.5). Non-canonical UV-B induced mismatches (not C → T or G → A) were filtered out using bcftools. Filtered mismatches were retained only if they were not present in other sequenced non-allelic lines without that phenotype. The resulting lists were filtered to retain only mismatches present in a CDS using bash scripting, then filtered manually in Interactive Genomics Viewer v 2.10 to remove mismatches which did not induce an amino acid change and which were misaligned or poor quality.

SNPs identified in the Mp*RTN4IP1L* gene in Mp*thar2*, Mp*thar4* and Mp*thar6* by the pipeline were confirmed by PCR amplification and Sanger sequencing using primers “GRT” (Table 3). Sanger sequencing was carried out by Source BioScience.

### Metabolomic analysis of pure and modified thaxtomin A in *M. polymorpha* thallus

Gemmae were grown on autoclaved cellophane (AA Packaging Limited, Preston, UK) on solid ½ Gamborg medium supplemented with 0.1 % DMSO for 14 days at 23 °C in 24 h light. Gemmalings were then transferred (by transfer of the cellophane disc) onto solid ½ Gamborg medium supplemented with either 0.1 % DMSO or 5 µM thaxtomin A and grown in these conditions for 2 days at 23 °C in 24 h light. Treated and untreated gemmalings were harvested by flash freezing in liquid nitrogen.

For thaxtomin A analysis, frozen tissue samples were ground in liquid nitrogen and homogenized in ice-cold extraction solvent (2:1:1 methanol:acetonitrile:H_2_O, v/v) for 2 min at 4 °C followed by incubation at -20 °C for one hour. Samples were centrifuged at full speed at 4 °C for 3 mins. Supernatants were collected and kept at -20 °C. Pellets were redissolved in ice cold 80 % (v/v) methanol and homogenized for 1 min at 4 °C followed by incubation at – 20 °C for one hour. Samples were centrifuged at full speed at 4 °C for 3 mins. Supernatants were collected and added to the supernatants from the first extraction step. Samples were incubated at -20 °C for 2 hours. Samples were then centrifuged at full speed at 4 °C for 10 mins. Supernatants were collected and shock frozen in liquid nitrogen. For coenzyme Q10 analysis, a chloroform/methanol extraction from frozen plant tissue was carried out.

Metabolite extracts were analyzed by the Metabolomics Facility at Vienna BioCenter Core Facilities (VBCF), member of the Vienna BioCenter (VBC), Austria and funded by the Austrian Federal Ministry of Education, Science & Research and the City of Vienna. For amino acid analysis, 100 µl of the extracts were dried down in a vacuum centrifuge and resolved in 100 µl of 0.1% formic acid in water. For each sample, 1 µl was injected onto a Kinetex (Phenomenex) C18 column (100 Å, 150 x 2.1 mm) connected with the respective guard column, and employing a 7-minute-long linear gradient from 99% A (1 % acetonitrile, 0.1 % formic acid in water) to 60% B (0.1 % formic acid in acetonitrile) at a flow rate of 80 µl/min. Detection and quantification was done by LC-MS/MS, employing the selected reaction monitoring (SRM) mode of a TSQ Altis mass spectrometer (Thermo Fisher Scientific), using the following transitions in the positive ion mode: 76 *m/z* to 30 *m/z* (glycine), 90 *m/z* to 44 *m/z* (alanine), 106 *m/z* to 60 *m/z* (serine), 116 *m/z* to 70 *m/z* (proline), 118 *m/z* to 72 *m/z* (valine), 120 *m/z* to 74 *m/z* (threonine), 132 *m/z* to 86 *m/z* (leucine and isoleucine),133 *m/z* to 74 *m/z* (asparagine), 134 *m/z* to 74 *m/z* (aspartic acid), 147 *m/z* to 84 *m/z* (lysine), 147 *m/z* to 130 *m/z* (glutamine), 148 *m/z* to 84 *m/z* (glutamic acid), 150 *m/z* to 133 *m/z* (methionine), 156 *m/z* to 110 *m/z* (histidine), 166 *m/z* to 133 *m/z* (phenylalanine), 175 *m/z* to 70 *m/z* (arginine), 176 *m/z* to 159 *m/z* (citrulline), 182 *m/z* to 36 *m/z* (tyrosine), 205 *m/z* to 188 *m/z* (tryptophan), 241 *m/z* to 74 *m/z* (cystine).

Thaxtomin A was analyzed by using a 4-minute-long linear gradient from 96% A to 90% B, directly analyzing the metabolite extract. Here, the transitions 439.1 *m/z* to 247.1 *m/z* and 439.1 *m/z* to 219.1 *m/z* were used in the positive ion mode. Quantification was done by external calibration using an authentic standard. The putative thaxtomin A metabolite (without the nitro-functionality) was discovered by analyzing the samples with precursor ion scanning for the dominant fragment ion (*m/z* 247.1) of thaxtomin A. Here only one compound was evident with a *m/z* of 394, sharing both dominant fragment ions (*m/z* 247.1 and *m/z* 219.1) with thaxtomin A). This metabolite was analyzed in the samples using the transitions 394.1 *m/z* to 247.1 *m/z* and 394.1 *m/z* to 219.1 *m/z*. Data interpretation was performed using TraceFinder (Thermo Fisher Scientific).

Coenzyme Q10 was analyzed after chloroform/methanol extraction from powdered frozen plant tissue. 100 µl of the lower phase was diluted with 50 µl methanol and 1 µl was directly injected onto a Kinetex (Phenomenex) C8 column (100 Å, 100 x 2.1 mm) and employing a 5-minute-long linear gradient from 90 % A (50 % acetonitrile, 49.9 % water, 0.1% formic acid, 10 mM ammonium formate) to 95% B (10 % acetonitrile 88% isopropanol, 1.9 % water, 0.1% formic acid, 10 mM ammonium formate) at a flow rate of 100 µl/min. Detection and quantification was done by LC-MS/MS, employing the selected reaction monitoring (SRM) mode of a TSQ Altis mass spectrometer (Thermo Fisher Scientific), using the transition 863.8 *m/z* to 197.1 *m/z* in the positive ion mode.

### DAB staining

3,3’-diaminobenzidine (DAB) (Sigma Aldrich: D8001) was used to prepare a DAB staining solution according to (46). Tak-1, Tak-2, Mp*rtn4ip1l^GE118^* and Mp*rtn4ip1l^GE149^*11-day old plants were incubated in 3 ml solution in 24 well plates and vacuum infiltrated for 5 minutes. The plates were covered in aluminium foil and incubated for 1.5 hours shaking at 100 rpm. After incubation, the staining solution was replaced by bleaching solution prepared according to (46). Plants were bleached for 2 hours. Bleached gemmae were imaged using a Keyence VHX-7000.

### Ferric-xylenol orange (FOX) assay

A modified FOX assay was carried out according to the protocol in (48). The FOX working solution was prepared as follows: 500 µM ammonium ferrous sulphate; 400 µM xylenol orange; 200 mM sorbitol; 25 mM H_2_SO_4_. Twelve-day old gemmalings of Tak-1, Tak-2, Mp*rtn4ip1l^GE118^* and Mp*rtn4ip1l^GE149^*were frozen and ground in liquid nitrogen and homogenized in 200 mM perchloric acid. Samples were centrifuged at 1000g for 5 minutes at 4 °C. 500 µl of the supernatant were mixed with 500 µl FOX working solution. Samples were incubated at room temperature in the dark for 30 minutes followed by quantification of the absorbance at 560 nm using an Ultrospec 3100 pro spectrophotometer. The concentration of H_2_O_2_ in each sample was calculated using a calibration curve of known concentrations of H_2_O_2_ quantified using the same protocol.

### Media

Modified Johnson’s medium was prepared as described in (75).

½ Gamborg medium:

| Component | Source | Final concentration |
| --- | --- | --- |
| Gamborg B5 medium basal salt mixture | Sigma Aldrich (G5768) | 1.5 g/l (w/v) |
| MES monohydrate | Applichem (A1074) | 0.5 g/l (w/v) |
| Sucrose | Applichem (A2211) | 1 % (w/v) |
| Agar | Applichem (A2111) | 0.8 % (w/v) |

pH adjusted to 5.5 using 1 M KOH

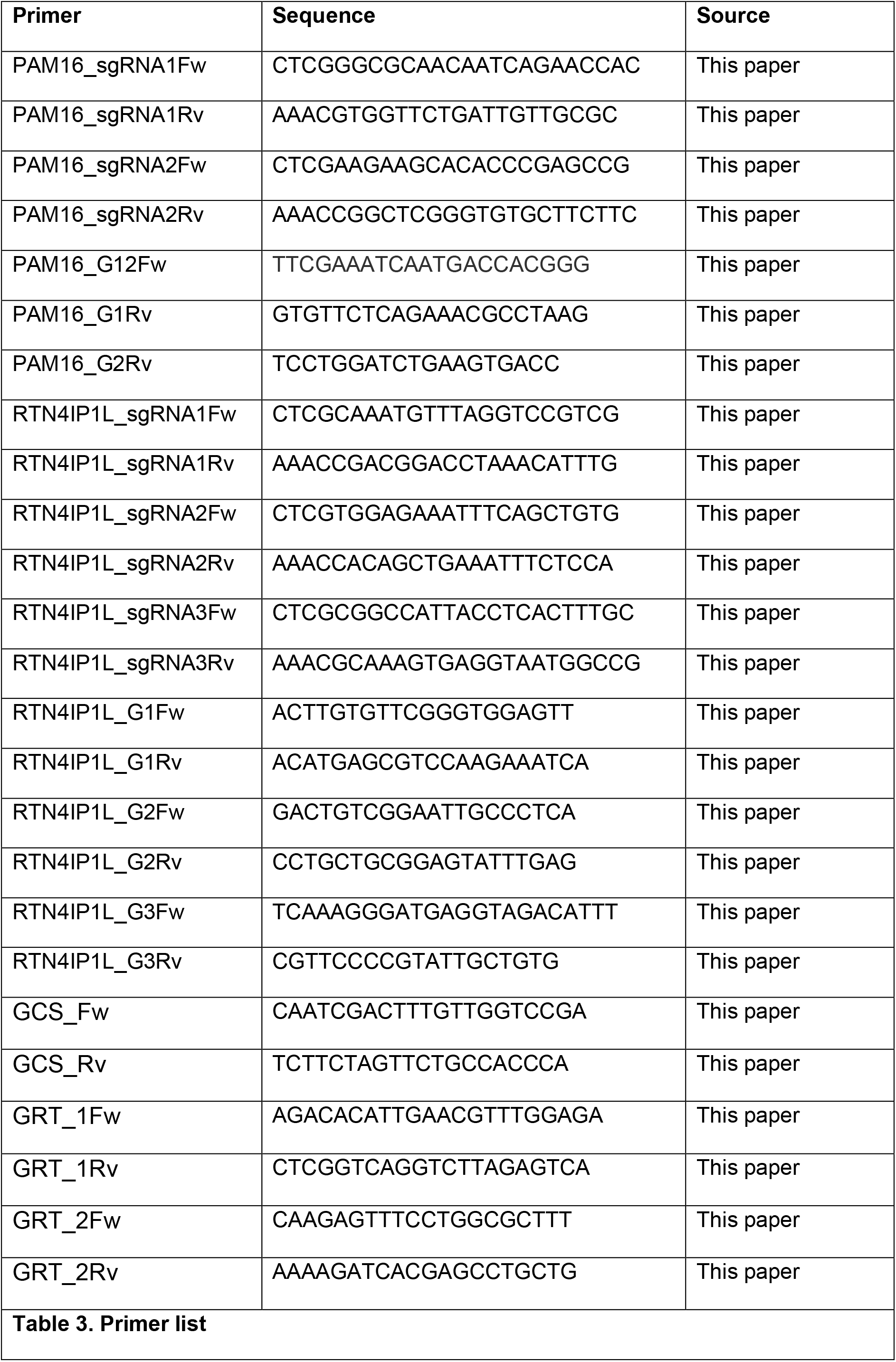

